# Human cytomegalovirus antagonizes SMC5/6 driven genome silencing via UL35 instituted proteasomal degradation of SLF2

**DOI:** 10.64898/2026.05.26.727865

**Authors:** Francesco Montagner, Myriam Scherer, Cora Stegmann, Friedrich Hahn, Claudia Ploil, Christina Sandra Einsiedler, Takahiro Maeki, Ramya Ramani, Nicole Andree-Busch, Melanie M. Brinkmann, Thomas Stamminger

## Abstract

Recent studies demonstrated that the structural maintenance of chromosomes complex 5/6 (SMC5/6) acts as an important antiviral restriction factor that silences episomal DNA. Consequently, viruses have evolved strategies to antagonize this defence system. Interestingly, it was shown that SMC5/6-mediated silencing of extrachromosomal DNA depends on its recruitment to PML nuclear bodies which is mediated by a protein termed SMC5/6 localization factor 2 (SLF2). Since human cytomegalovirus (HCMV) dissociates PML nuclear bodies (PML-NBs) during the first hours of infection, we asked for the fate of SMC5/6 components during the HCMV replicative cycle. We first investigated the expression levels of SMC5/6 core components at the onset of HCMV infection and found that they were not downregulated by HCMV. Instead, we observed a distinct decrease of SLF2 protein levels which correlated with a complete dispersal of the SMC5/6 complex from PML-NBs. This was also observed after infection with UV inactivated HCMV and could not be blocked by cycloheximide treatment suggesting the involvement of an imported structural component of the virion. While we could exclude a role of the HCMV tegument protein pp71 (UL82), a targeted screen identified the tegument protein UL35 as being required and sufficient for proteasomal degradation of SLF2. In line with these results, SLF2 levels remained stable after infection with a recombinant HCMV harbouring a stop codon within the UL35 gene region and this correlated with a defect in the onset of HCMV immediate early gene expression. Furthermore, we observed an enhanced recruitment of SMC5/6 to PML-NBs and an overall increase in the association of parental viral genomes with SMC5/6 containing PML-NBs in the absence of UL35. Congruently, siRNA mediated depletion of SLF2 resulted in a reversal of this phenotype. This strongly suggests that UL35 serves as a novel viral SLF2 antagonist to prevent the initial silencing of incoming viral genomes by SMC5/6.

**Author summary:** Human cytomegalovirus (HCMV) is an important human pathogen which establishes a life-long persistent infection. Previous research revealed that host intrinsic defense mechanisms can silence HCMV gene expression and thereby foster latency. However, during lytic infection, viral effector proteins like the structural protein pp71 (UL82) and the viral immediate-early protein IE1 act as antagonists of this defense. Here, we report on the identification of a novel antagonist of host-mediated viral gene silencing which targets the SMC5/6 complex. SMC5/6 belongs to a highly conserved family of protein complexes that act as loop extrusion machines to facilitate the folding of chromatin. While SMC5/6 core factors were not attacked by HCMV, we found that the SMC5/6 associated protein SLF2 was degraded by the viral tegument protein UL35, that is imported into infected cells. This abrogates the accumulation of SMC5/6 at PML nuclear bodies which compromises the entrapment of viral genomes in these subnuclear structures. Since SLF2 is also important for genome stability and pathogenic variants of SLF2 cause the neurodevelopmental disease Atelis syndrome, we speculate that depletion of this factor via a viral tegument protein may also contribute to DNA damage induced during congenital HCMV infection.

## Introduction

Immune sensing of foreign nucleic acids constitutes a hallmark of virus detection [1]. This becomes particularly challenging when pathogenic DNA inside the cell nucleus needs to be recognized driving the evolution of functional intersections between DNA damage response (DDR) and antiviral defense mechanisms [2]. Recent studies demonstrated that the structural maintenance of chromosomes (SMC) complex 5/6 acts as an emerging antiviral restriction factor with the capacity to silence episomal DNA [3, 4]. SMC complexes serve as important genome regulators which are highly conserved across species and include SMC1/3 (cohesin), SMC2/4 (condensin) and SMC5/6 [5]. They form ring-shaped molecular machines with ATPase activity that allows folding of genomic DNA into loops [5]. This plays essential roles in diverse biological processes such as the separation of replicated DNA molecules or the facilitation of enhancer-promoter interactions [6]. SMC5/6 differs from other SMC complexes with regard to a more local and targeted action on chromosomes essential for DNA repair and genome stability [7]. It is also unique concerning the subunit composition of its core holoenzyme comprising the SMC5/6 heterodimer and additional proteins termed non-SMC elements (NSMCE) 1, 3 and 4a [7] (Fig. 1A). Furthermore, it associates with cellular proteins like SMC5/6 localization factor 1 (SLF1) and 2 (SLF2) which recruit the complex to stalled replication forks [8]. In contrast, a different subcomplex containing SLF2 has recently been shown to mediate the SMC5/6 driven silencing of episomal DNA [9]. Thus, SLF2 regulatory subcomplexes appear to direct distinct roles of SMC5/6 on chromosomal versus episomal DNA [9] (Fig. 1A).

**Fig. 1:**
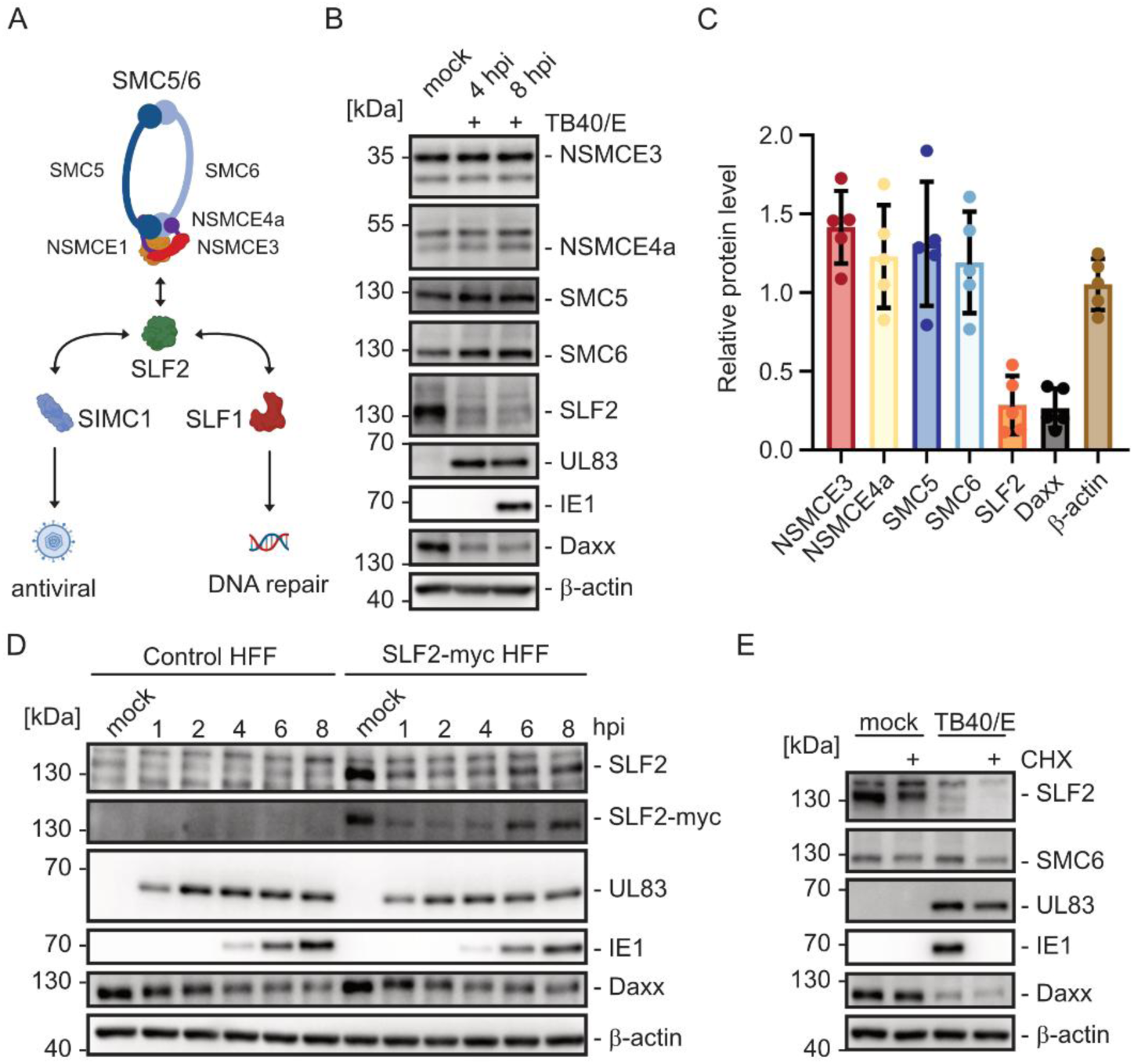
Downregulation of SLF2 but not of SMC5/6 core components during the first hours after HCMV infection. (A) Schematic representation of SMC5/6 core components and the role of SLF2, SIMC1 and SLF1 to direct SMC5/6 either to antiviral or DNA repair pathways. (B-C) Investigation of SMC5/6 protein levels upon HCMV infection. HFF cells were either not infected (mock) or infected at a multiplicity of 3 (MOI 3) for 4 or 8 hours (hpi). (B) Whole cell lysates were harvested for immunoblotting analysis of the levels of the core components of SMC5/6 and SLF2. Cellular Daxx and the viral UL83 and IE1 proteins were included as controls, as well as β-actin as loading control. (C) Five independent biological replicates of (B) were performed and the average fold change of protein levels was plotted +/- standard deviation (SD). (D) HCMV-mediated downregulation of SLF2 after overexpression of myc-tagged SLF2 in HFF. Whole HFF lysates were collected from control cells and cells overexpressing SLF2-myc that were either mock infected or infected with HCMV TB40/E (MOI 3) for 1 to 8 hours as indicated. SLF2 expression was either detected with SLF2 specific polyclonal or myc-specific monoclonal antibodies. Cellular β-actin served as loading control. One representative result out of three biological replicates is shown. (E) HFF cells, either mock infected or infected with HCMV TB40/E (MOI 1) were left untreated or were treated with 150 µg/ml of cycloheximide (CHX). At 8 hpi, whole cell lysates were collected and SLF2 protein levels analyzed. The viral IE1 protein served as control for successful protein synthesis inhibition; β-actin was used as loading control. One representative experiment out of three biological replicates is shown.

Episomal genomes play essential roles in the life cycle of several viruses. For instance, herpesviruses are thought to circularize their linear DNA genomes immediately upon entry into the cell nucleus and might thus serve as targets for SMC5/6 repression [10]. Consequently, recent evidence suggests the existence of viral antagonistic mechanisms to evade SMC5/6 as an intrinsic immune sensor and restriction factor. For Epstein-Barr virus, the major tegument protein BNRF1 has been shown to degrade the SMC5/6 core components via a proteasomal mechanism [11]. There is also evidence that SMC5/6 recognizes the genomes of another γ-herpesvirus, Kaposi sarcoma-associated herpesvirus (KSHV), to inhibit its replication via generation of a repressive chromatin structure [12]. This is counteracted by the KSHV protein RTA and its homologs in monkey and mouse γ-herpesviruses which act as E3 ubiquitin ligases to target the core components SMC5 and SMC6 for degradation [12].

In contrast, SMC5/6 antagonizing proteins have not been identified within the subfamilies of α- and β-herpesviruses, yet. Human cytomegalovirus, an important pathogen for immunosuppressed patients, is the prototype of the β-herpesviruses which are characterized by their prolonged replication cycle [13]. Previous studies demonstrated that silencing of incoming HCMV genomes is mediated by a subnuclear structure termed promyelocytic leukemia (PML) nuclear bodies (PML-NBs) which are biomolecular condensates formed by multiple proteins including the core components PML, Sp100, Daxx and ATRX [14–16]. DNA labeling revealed that invading HCMV genomes are entrapped inside PML-NBs and remain stably associated with PML cages in a transcriptionally repressed state in the absence of viral antagonization [17]. However, during lytic infection HCMV successfully counteracts this cellular repression mechanism via the action of two viral effector proteins: (i) the tegument protein pp71 (UL82) targets the chromatin remodeling complex Daxx/ATRX to dissociate ATRX from PML-NBs and to induce the proteasomal degradation of Daxx [18, 19]; (ii) the major immediate early protein IE1 directly interacts with the TRIM family member PML via coiled-coil interactions [20, 21]. This inhibits the de novo SUMOylation of PML which dissociates PML-NBs thus interfering with the entrapment of viral DNA [17, 22–24].

Here, we identify the first SMC5/6 antagonistic protein of β-herpesviruses. In contrast to γ-herpesviruses, which were shown to deplete SMC5/6 core components, we observed the selective proteasomal degradation of the regulatory subcomplex protein SLF2 instituted by the HCMV tegument protein UL35. This abrogated the localization of SMC5/6 complexes at PML-NBs. The delocalization of SMC5/6 from PML-NBs upon SLF2 degradation suggests a novel mechanism for how herpesviruses prevent genome silencing by intrinsic cellular defenses.

## Results

### HCMV induces a downregulation of SLF2 but not of SMC5/6 core components immediately after infection

SMC5/6 is unique among the structural maintenance of chromosomes (SMC) complexes in its broad ability to silence the transcription of extrachromosomal circular DNA elements which appears to restrict the replication of several distinct viruses [3, 25]. Since recent studies suggested that viruses may antagonize this antiviral activity via the degradation of SMC5/6 complex components, we asked the question whether this is also true for human cytomegalovirus (HCMV) [3]. To investigate this, primary human foreskin fibroblasts (HFF) were infected with HCMV strain TB40/E and cell lysates harvested at 4 and 8 hours post infection (hpi). Immunoblotting with antibodies against SMC5, SMC6, NSMCE3 and NSMCE4a showed that the abundance of these SMC5/6 core proteins was not significantly modulated upon HCMV infection (Figure 1A, B). However, we observed a decrease in SMC5/6 localization factor 2 (SLF2) protein abundance with kinetics similar to the well characterized HCMV restriction factor Daxx which undergoes proteasomal degradation prior to the onset of HCMV immediate early (IE) gene expression (Figure 1B, C) [18]. To further confirm a specific modulation of SLF2 abundance by HCMV, we established HFF cells overexpressing myc-tagged SLF2. Using both myc-specific monoclonal and SLF2-specific polyclonal antibodies we found that HCMV infection resulted in a transient decrease of SLF2 abundance immediately upon and during the first hpi (Figure 1D). Of note, downregulation of SLF2 was also detected after infection of the retinal pigment epithelial cell line ARPE-19 suggesting a cell-type independent mechanism (S1 Figure). Furthermore, although treatment of cells with the protein synthesis inhibitor cycloheximide slightly affected the abundance of SLF2 in non-infected cells, it did not prevent the strong downmodulation of SLF2 after infection with HCMV (Figure 1 E). Since cycloheximide blocks the onset of de novo viral protein synthesis, but does not affect viral structural proteins that are imported with viral particles, viral immediate-early proteins are most probably not required to regulate SLF2 abundance (Figure 1E). Thus, we hypothesize that either viral entry or the import of a structural protein is necessary for the infection-induced decrease of SLF2 protein expression.

### HCMV infection dissociates SMC5/6 from PML-NBs

In a cell culture infection model with hepatitis B virus it has been shown that SLF2 is necessary to recruit the SMC5/6 complex to PML nuclear bodies [26]. Consequently, we wanted to clarify whether HCMV infection affects the subnuclear localization of SMC5/6 in HFF. First, SMC5/6 antibodies were evaluated concerning their suitability for immunolocalization experiments. Labeling of HFF cells with antibodies against SMC6 and NSCME3 revealed a dot-like pattern resembling the subnuclear accumulation of PML-NBs (Figure 2A). To evaluate the colocalization of individual proteins, we established a method based on determination of Pearson’s correlation coefficient (PCC) (S2 Figure). Since a high degree of colocalization between SMC6 and NSMCE3 was observed, we assume that these dot-like structures represent the intact SMC5/6 core complex possibly localizing at PML-NBs (Figure 2B). To investigate the role of SLF2 for this dot-like distribution pattern of SMC5/6, we established conditions for siRNA-mediated knockdown. After having confirmed the successful depletion of SLF2 following transfection of specific siRNAs by Western blot analysis (Figure 2C), its effect on the subnuclear accumulation of SMC5/6 was investigated. Using NSMCE3 as a marker protein we observed a distinct colocalization between the SMC5/6 complex and PML-NBs in non-transfected HFFs and cells transfected with a control siRNA (siC) (Figure 2D, E; siC). In contrast, siRNA-mediated depletion of SLF2 resulted in a highly significant dissociation of SMC5/6 from PML-NBs indicating a requirement of SLF2 for SMC5/6 localization at PML-NBs (Figure 2D, E; siSLF2). Due to the observed downregulation of SLF2 by HCMV, we then performed infection experiments to investigate the colocalization of SMC5/6 with PML-NBs at the onset of viral gene expression. At 8 hpi, when IE1-induced dispersal of PML-NBs was still incomplete, a highly significant dispersal of SMC5/6 from PML-NBs was visible (Figure 2F, G). In addition, infection was performed in the presence of CHX to prevent IE1 protein expression. Although CHX treatment slightly reduced the association between SMC5/6 and PML-NBs in mock infected cells, it did not reverse the strong infection-induced dispersal, further confirming that viral immediate early proteins are dispensable for this effect (Fig. 2F, G, CHX).

**Fig. 2:**
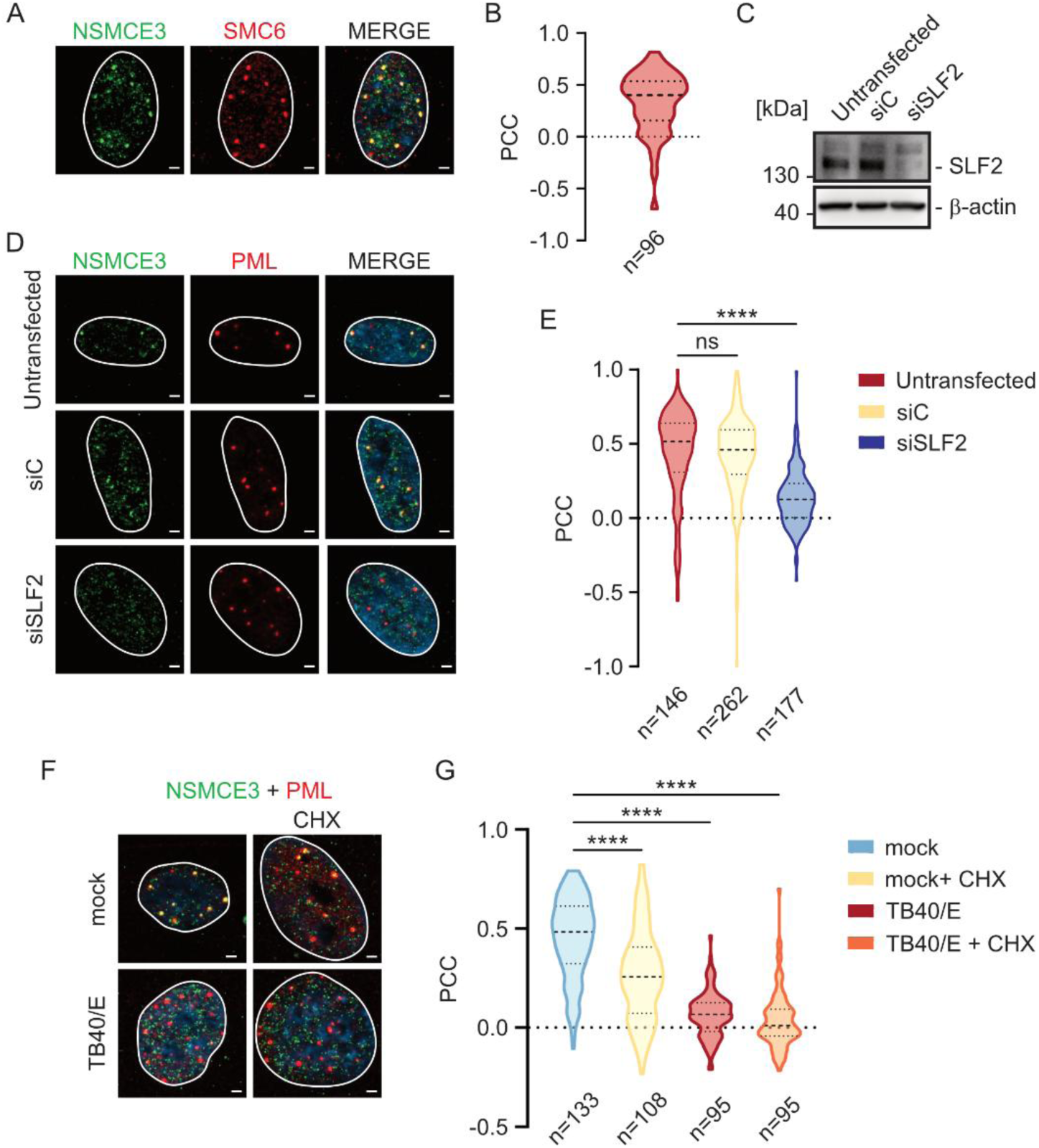
Effect of SLF2 depletion and HCMV infection on the association of SMC5/6 with PML-NBs. (A-B) Subnuclear localization of SMC5/6 core components in HFFs. (A) Indirect immunofluorescence staining of NSMCE3 and SMC6 showing the colocalization of both proteins in dot-like subnuclear accumulations. Scale bar: 2 µm. (B) Colocalization analysis of SMC6 and NSMCE3. A total of 96 nuclei from three independent biological replicates were analysed using the Zen Blue colocalization module. The Pearson’s correlation coefficients (PCC) between SMC6 and NSMCE3 were calculated in an area restricted to the foci of SMC6 and visualized in a violin plot. The median, the first and third quartile are indicated. (C-E) Colocalization of the SMC5/6 core complex with PML-NBs depends on SLF2. (C) Untransfected HFFs or HFFs transfected with a control siRNA (siC) or siRNAs against SLF2 (siSLF2) were either harvested or immunolabeled to detect PML and NSMCE3. (C) Western blot analysis to confirm the knockdown of SLF2; β-actin served as loading control. (D) Indirect immunofluorescence analysis of PML and NSCME3 in untransfected or siRNA-transfected HFFs. (E) Quantification of the colocalization of PML and NSMCE3 through the PCC of a region of interest (ROI) surrounding the PML foci. A total of more than 140 nuclei from two biological replicates for the untransfected condition and three biological replicates for siC and siSLF2 conditions were analysed and the resulting PCCs displayed in a violin plot. The median, the first and third quartile are indicated. Statistical differences were calculated with one-way ANOVA (Kruskal-Wallis test): n.s.: non-significant, **** p< 0.0001. (F-G) Effect of HCMV infection on the colocalization of SMC5/6 with PML-NBs. (F) Indirect immunofluorescence analysis of the colocalization of NSMCE3 with PML-NBs upon HCMV infection. HFF cells were infected for 8 hours in absence or in presence of 150 µg/ml of cycloheximide (CHX) before they were harvested for immunofluorescence staining of PML and NSMCE3. A single cell with PML-NBs still intact was chosen as representant of the infected condition. Scale bar: 2 µm. (G) The PCCs of more than 90 cells from three biological replicates were quantified and the population distribution visualized in a violin plot. The median, the first and third quartile are indicated. In the infected conditions, only cells with intact PML were included in the analysis. Statistical differences were calculated with one-way ANOVA (Kruskal-Wallis test): ****p < 0.0001.

### A viral structural protein degrades SLF2 in a proteasome-dependent manner

To further understand how HCMV downregulates SLF2 we first performed infection experiments with an attachment competent but entry-deficient HCMV exhibiting the insertion mutation A332 in UL128 (TB40/E UL128insA332, kindly provided by C. Sinzger, Ulm) [27].

As shown in Fig. 3A, neither SLF2 nor Daxx were downregulated upon infection with TB40/E-UL128insA332. In contrast, protein levels of the tegument protein pp65 (UL83) were diminished confirming the defect in viral entry (Figure 3A). We conclude that viral attachment is not sufficient to trigger the decrease of SLF2 protein levels. Next, we performed infection experiments with UV-inactivated viral inocula to investigate whether imported viral proteins are involved (Figure 3B-D). Infection with a UV-inactivated TB40/E expressing IE2 fused to the autofluorescent EYFP resulted in both SLF2 and Daxx downregulation (Figure 3B). This correlated with a highly significant dispersal of SMC5/6 from PML-NBs after infection with UV-inactivated virus (Figure 3C, D). These results indicate that a viral structural protein imported by infection, but not de novo viral transcription, may be responsible for modulation of SLF2 protein levels. To investigate whether SLF2 downregulation can be blocked by inhibitors, we tested the effect of the proteasome inhibitor MG132, the ubiquitin-activating enzyme E1-inhibitor PYR-41 and the neddylation inhibitor MLN4924. Western blot experiments demonstrated that treatment of cells with MG132 resulted in a strong rescue of both SLF2 and Daxx protein levels suggesting that, similar to Daxx, SLF2 undergoes proteasomal degradation upon infection (Fig. 3E) [18]. Dispersal of SMC5/6 from PML-NBs was also inhibited by MG132 although the rescue was not complete (Fig. 3 E, F). This might be due to a negative effect of MG132 on the SMC5/6 association with PML-NBs already observed in mock infected cells (Fig. 3F). In summary, these results indicate that a viral structural protein degrades SLF2 in a proteasome-dependent manner.

**Fig. 3:**
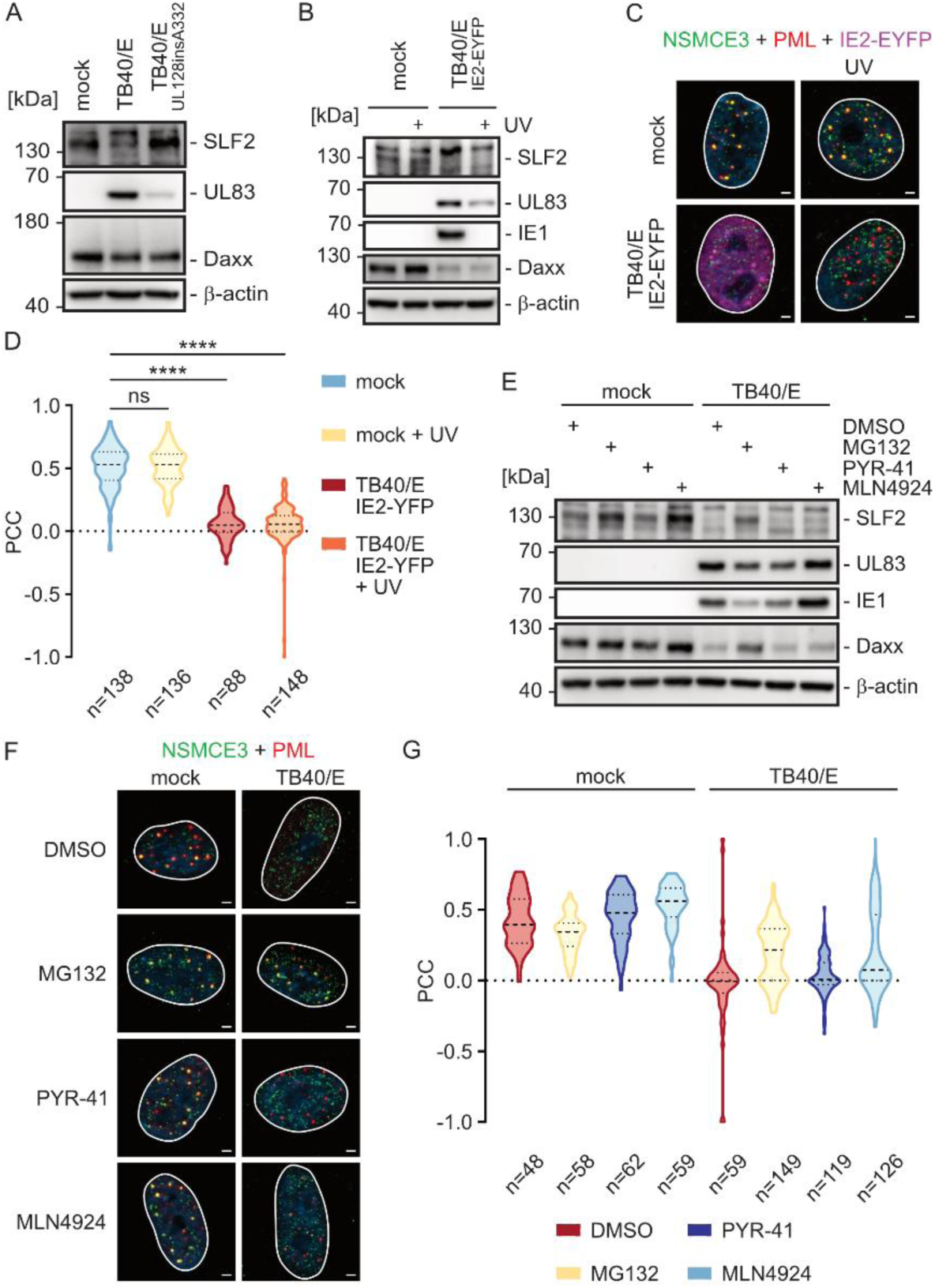
Evidence for a structural protein-driven proteasomal degradation of SLF2. (A) Requirement of viral entry for SLF2 degradation. ARPE-19 cells were infected for 4 hours (MOI 1) either with an entry proficient strain of TB40/E (TB40/E BAC_KL17_ UL32-EGFP UL100-mCherry) or with the entry-deficient strain TB40/E UL128insA332 (TB40/E BAC_KL17_ UL128insA332 UL32-EGFP UL100-mCherry) or were left uninfected (mock). Whole cell lysates were analysed by immunoblotting, and the SLF2 protein levels compared to mock cells. Viral UL83 was detected as control for viral entry and β-actin as loading control. Three biological replicates were performed. (B-D) Evidence for SLF2 degradation by a viral structural protein. (B) HFF cells were either left uninfected (mock) or were infected for 8 hours (MOI 1) with TB40/E IE2-EYFP. Where indicated, supernatants were UV-inactivated and added to the respective condition. The expression levels of SLF2 compared by western blotting. UV inactivation of the viral inoculum was controlled via the detection of IE1 expression. The experiment was performed in three biological replicates. (C) Indirect immunofluorescence analysis to detect the colocalization of PML and NSMCE3 either after mock infection or after infection of HFF with TB40/E IE2-EYFP. UV: UV treatment of supernatants. Scale bar: 2 µm. (D) Colocalization analysis of NSMCE3 and PML via quantification of PCCs. The analysis was restricted to the PML signals in the cell nuclei. More than 90 cells were analysed and the violin plot shows results of two independent biological replicates for the TB40/E IE2-EYFP sample and of three biological replicates for the other tested conditions. The median, the first and the third quartile are indicated. One-way ANOVA (Kruskal-Wallis test): n.s.: not significant, **** p< 0.0001. (E-F) SLF2 is degraded in a proteasome-dependent manner. (E) Western blot analysis of HFFs that were either mock infected or infected with TB40/E (MOI 1) for 8 hours. As indicated, cells were treated with 10 µM of MG132, 5 µM of PYR-41, 10 µM of MLN4924 or DMSO. SLF2, UL83, IE1 and Daxx were detected using specific antibodies; β-actin served as a loading control. (F) Indirect immunofluorescence analysis to detect the colocalization of NSMCE3 and PML. Infection and inhibitor treatment were performed as described for panel E. Scale bar: 2 µm. (F) Colocalization analysis of PML and NSMCE3 via quantification of PCCs. The violin plot summarizes the results of three independent biological replicates. The median, the first and the third quartile are indicated.

### The tegument protein pp71 (UL82) and the immediate early protein IE1 are not required for SLF2 degradation

Our experiments revealed that both SLF2 and Daxx are downmodulated via proteasomal degradation immediately upon HCMV infection (Fig. 3E). Thus, it was suggestive to ask whether the tegument protein pp71 (UL82), known to affect Daxx protein levels, also regulates SLF2 [18]. To answer this, we constructed a recombinant HCMV TB40/E with stop codons within UL82 thereby abrogating pp71 (UL82) expression (S3 Figure). In addition, an IE1 deletion virus was used to exclude PML-NB dispersal as determinant of SLF2 degradation [17]. Infection with TB40/E UL82stop resulted in efficient SLF2 degradation which was also observed for infection with the IE1 deletion virus (Fig. 4A). Furthermore, both viruses lacking either IE1 or pp71 expression were able to disperse SMC5/6 from PML-NBs (Fig. 4B, C). This finally demonstrated that neither the tegument protein pp71 nor the immediate early protein IE1 affect SLF2 and/or SMC5/6.

**Fig. 4:**
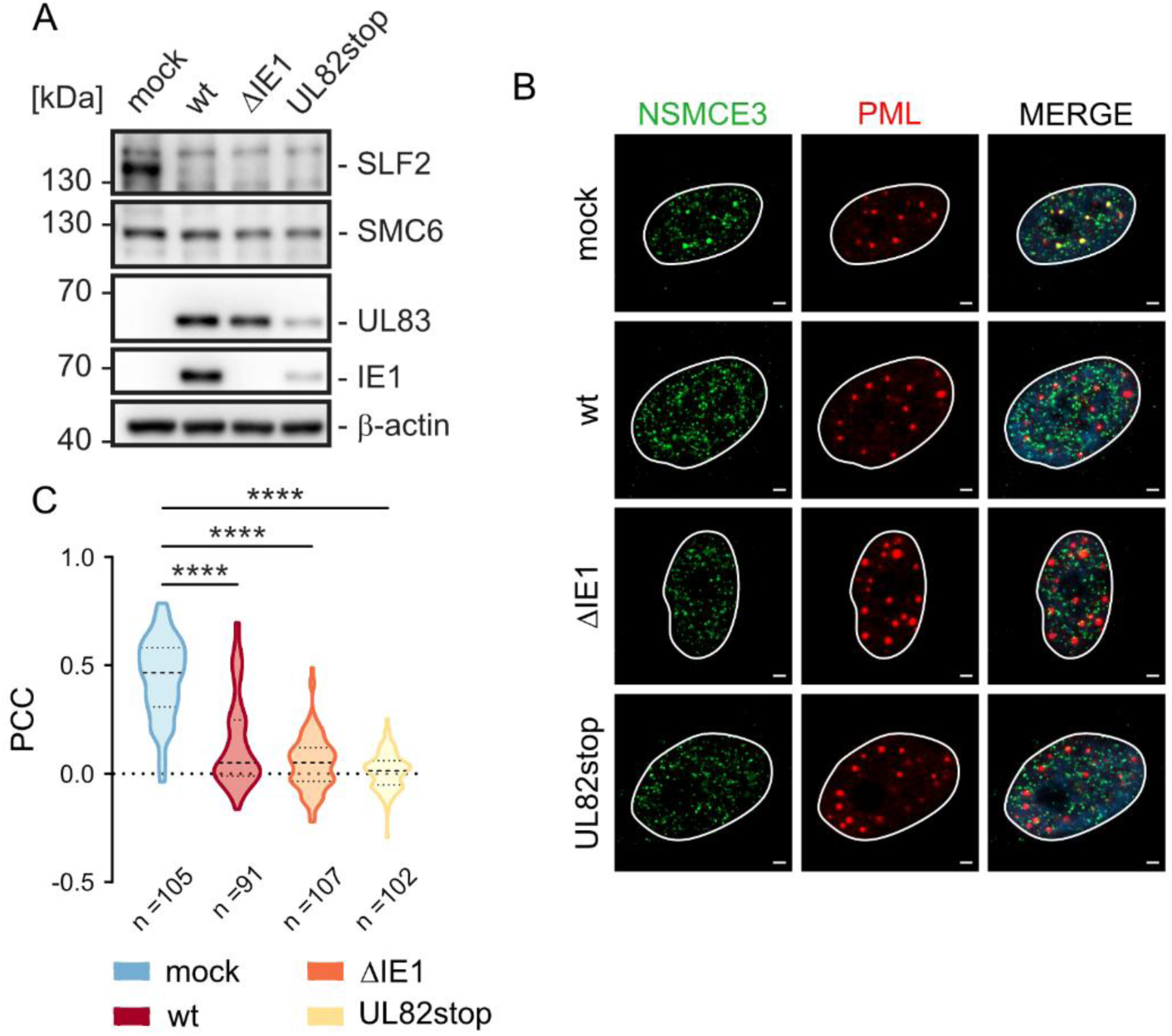
SLF2 degradation does not require IE1 or pp71 (UL82). (A-C) HFF were either mock infected or infected with wildtype TB40/E (wt), IE1-deleted TB40/E (ΔIE1), or a TB40/E harbouring a stop codon in UL82 (UL82stop) at an MOI of 1. After 8 hours, the cells were harvested and either whole cell lysates were prepared for western blot analysis or fixed for subsequent immunofluorescence analyses. (A) Western blot analysis to detect SLF2, SMC6, UL83 and IE1 protein levels; β-actin served as a loading control. (B) Immunofluorescence analysis to detect the colocalization of NSMCE3 and PML. Scale bar: 2 µm. (C) Colocalization analysis of PML and NSMCE3 via quantification of PCCs. The violin plot summarizes the results derived from three independent biological replicates. The median, the first and third quartile are indicated. Statistical differences were tested for with One-way ANOVA (Kruskal-Wallis test); ****, p<0.0001.

### A targeted screen identifies the tegument protein UL35 as the viral factor mediating SLF2 degradation

To identify the viral factor mediating SLF2 degradation, we tested several candidate proteins in a targeted screening approach. Plasmids encoding the respective viral proteins together with an expression vector for SLF2 were used to co-transfect ARPE-19 cells followed by detection of proteins via indirect immunofluorescence analysis (Fig. 5A). Subsequently, the cells were scored for the expression of SLF2, candidate protein or the co-expression of both (Fig. 5B). This strategy revealed that upon co-transfection of an expression plasmid for the tegument protein UL35, cells expressing both SLF2 and UL35 were missing (Fig. 5A, B). To further confirm that UL35 is responsible for the degradation of SLF2, HFFs with dox-inducible expression of HA-tagged UL35 were analysed [28]. In Western blot experiments, we observed that dox induction of UL35 protein expression correlated with significantly reduced levels of SLF2 while the core component SMC6 was not affected (Fig. 5C). Furthermore, immunolocalization studies revealed a distinct, dot-like subnuclear distribution of SLF2 in control cells (Fig. 5D). In contrast, expression of UL35 abrogated the detection of SLF2 dots (Fig. 5D, E). These results strongly suggest UL35 as the viral factor responsible for degradation of SLF2.

**Fig. 5:**
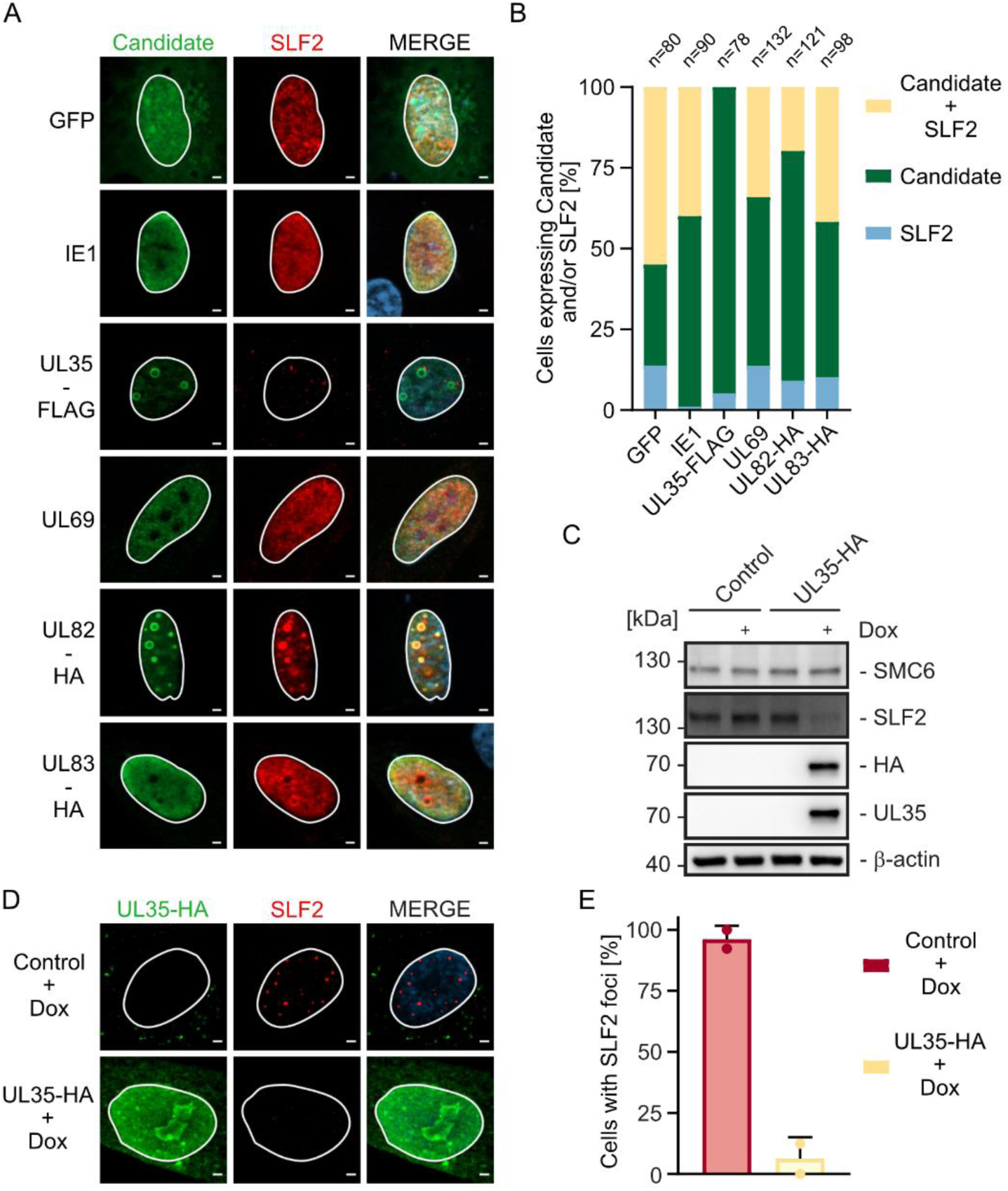
The tegument protein UL35 is sufficient for SLF2 degradation. (A-B) Targeted screening for SLF2 degradation in ARPE-19 cells. (A) ARPE-19 cells were co-transfected with expression vectors for SLF2 and candidate viral effector proteins. Vectors encoding GFP and IE1 were included as controls. The cells were harvested at 24 hours after transfection for indirect immunofluorescence staining. Scale bar: 2 µm. (B) Quantification of cells positive for candidate and/or SLF2 protein. The expression patterns of more than 70 cells were counted from three independent biological replicates and displayed as percentage (%) of cells expressing SLF2, the candidate protein or both. (C-D) SLF2 degradation in HFFs with doxycycline-inducible expression of HA-tagged UL35. Control and UL35-HA inducible cells were stimulated with 1µg/ml of doxycycline (dox) for 24 hours. (C) Western blot analysis to detect endogenous SLF2 and SMC6 after doxycycline-induction of UL35-HA expression; β-actin served as a loading control. (D) Immunofluorescence analysis of the subcellular localization of SLF2 after induction of UL35-HA expression by doxycycline. UL35 expression was detected using a monoclonal antibody against the HA-tag (HA); SLF2 was detected with a SLF2 specific polyconal antibody (SLF2). Scale bar: 2 µm. (E) Quantification of the number of cells with SLF2 foci in the absence or presence of UL35-HA. Two independent biological replicates were performed and >70 cells were analysed. The means of the % were calculated +/- SD.

### UL35 but not UL35a mediates the degradation of SLF2

Previous studies showed that the UL35 gene region encodes two proteins, UL35 and UL35a, that are expressed at different times of the replication cycle and also differ in function [29]. The larger protein, UL35, is only produced at late times of the replication cycle [29]. However, it is incorporated into the tegument of virions and therefore imported into newly infected cells [30]. UL35a is synthesized via the control of an internal promoter with early-late kinetics, giving rise to a smaller protein which comprises the C-terminal part of UL35 [29]. While UL35 has the ability to form PML-altering nuclear bodies, UL35a lacks this activity [31]. To differentiate whether UL35 or UL35a mediates SLF2 degradation, we co-transfected ARPE-19 cells with expression plasmids for FLAG-tagged UL35 or UL35a and SLF2. In addition, plasmids for the HIV-1 accessory protein Vpr (Vpr) and the HCMV tegument protein pp65 (UL83) were included as positive and negative controls. Immunodetection and subsequent scoring for co-expression revealed that both Vpr and UL35, however not UL35a and UL83, abrogated the expression of SLF2 (Fig. 6A, B). To confirm this, HFFs with dox-inducible expression of either FLAG-tagged UL35 or UL35a were generated. As shown in Fig. 6C, only UL35, but not UL35a, was able to deplete endogenous SLF2 (Fig. 6C). This could be inhibited by the addition of MG132, confirming that the isolated expression of UL35 is sufficient to induce SLF2 degradation via the proteasome (Fig. 6D). Finally, only upon expression of UL35 we observed the previously described alteration of PML-NBs and the disappearance of SLF2 foci in immunolocalization experiments (Fig. 6E) [31].

**Fig. 6:**
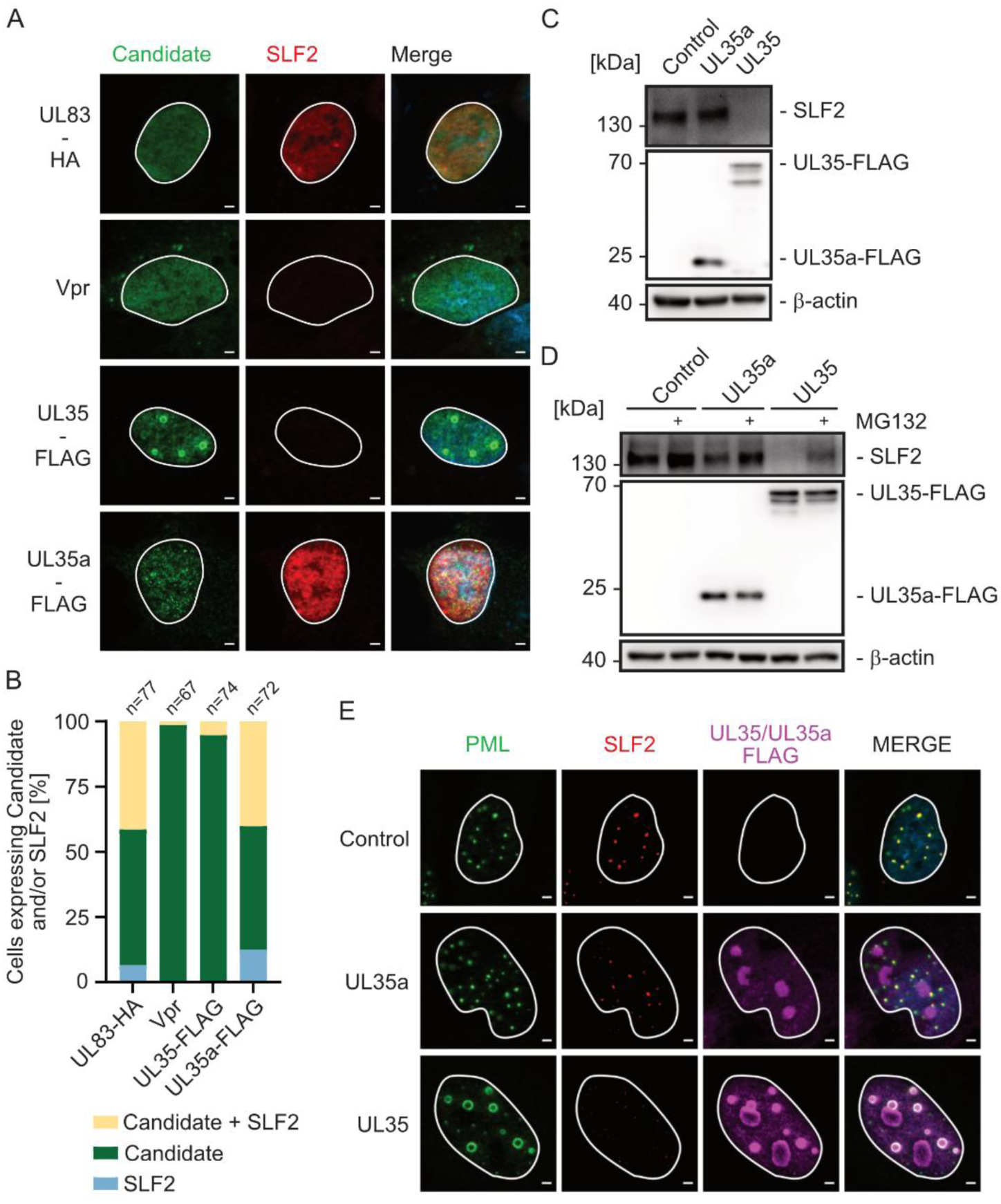
UL35 but not UL35a mediates the degradation of SLF2. (A-B) Coexpression of UL35a and SLF2 are not mutually exclusive. ARPE-19 cells were cotransfected with an expression plasmid encoding SLF2 together with plasmids for FLAG-UL35, FLAG-UL35a, UL83 or Vpr. (A) Indirect immunofluorescence analysis, performed at 24 hours post transfection, to detect SLF2 and the respective candidate proteins using a SLF2 specific polyconal antibody together with antibodies directed against UL83, Vpr, or the FLAG-tag. Scale bar: 2 µm. (B) Quantification of cells positive for candidate and/or SLF2 protein. The expression patterns of more than 60 cells were counted from three independent biological replicates and displayed as percentage (%) of cells expressing SLF2, the candidate protein or both. (C-E) Dox-inducible expression of UL35a is not sufficient for SLF2 degradation. Control cells or HFFs with doxycycline-inducible expression of either UL35a-FLAG or UL35-FLAG were incubated with 500 ng/ml of doxycycline for 24 hours followed by either the preparation of cell lysates for western blotting or the fixation of cells for subsequent immunofluorescence analysis. (C and D) Western blot analysis to detect endogenous SLF2 as well as UL35 and UL35a using a FLAG-specific antibody; β-actin was used as a loading control. Panel D includes the analysis of cell lysates that were harvested after incubation with MG132 (10 µM) or DMSO to investigate the rescue of SLF2 via proteasomal inhibition. Three independent biological replicates were performed. (E) Indirect immunofluorescence analysis of the subcellular localization of PML and SLF2 upon expression of either UL35 or UL35a. Three independent biological replicates were performed. Scale bar: 2 µm.

### Abrogation of SLF2 degradation correlates with a defect to initiate immediate early gene expression

So far, we showed that SLF2 undergoes proteasomal degradation upon HCMV infection, and expression of UL35 was sufficient to induce SLF2 depletion, which annihilates the colocalization of SMC5/6 with PML-NBs. Since SMC5/6 is supposed to repress episomal viral transcription we were interested in the phenotype of an HCMV UL35 null mutant concerning the onset of immediate early gene expression. For this, we used an HCMV TB40/E mutant harboring a stop codon at nucleotide position 228 of the UL35 ORF which abrogates UL35, but not UL35a expression [28]. This was important because previous studies have shown that deletion of the entire UL35 or of the UL35a coding region results in a severe growth deficit due to a defect in virus assembly and particle formation [32, 33]. First, we investigated whether deletion of the UL35 protein interfered with SLF2 degradation in the context of viral infection. Indeed, when HFFs were infected with TB40/E UL35stop, titrated for equivalent IE1 expression, SLF2 protein levels were similar to mock while Daxx protein levels were diminished (Fig. 7A, B). In contrast, infection with TB40/E UL82stop resulted in diminished SLF2 but preserved Daxx protein levels indicating a selective degradation of either SLF2 or Daxx by UL35 or pp71 (UL82), respectively (Fig. 7A). After having demonstrated a defect in SLF2 degradation we wanted to know whether this correlates with a reduced infectivity of TB40/E UL35stop. Since an effect on the IE phase was expected, it was important to standardize the inoculum by determination of viral DNA imported into infected cell nuclei. We used qPCR to standardize the inocula to equal genome copies for both wildtype and mutant viruses. HFFs were then infected with equivalent inocula (MOIs of 0.05 and 0.005) of either TB40/E wt or UL35stop. At 24 hpi cells were fixed, stained for IE1 and the number of IE1 positive cells, representing cells with successful initiation of viral of immediate-early gene expression, was determined. This revealed a significantly lower number of IE1 positive cells after infection with TB40/E UL35stop (Fig. 7C). To corroborate this, we decided to analyze viral transcript levels at 8 hpi. As shown in Fig. 7D, both IE1 and IE2 specific transcripts were reduced indicating that the lack of SLF2 degradation correlates with a defect in the onset of viral immediate early gene transcription.

**Fig. 7:**
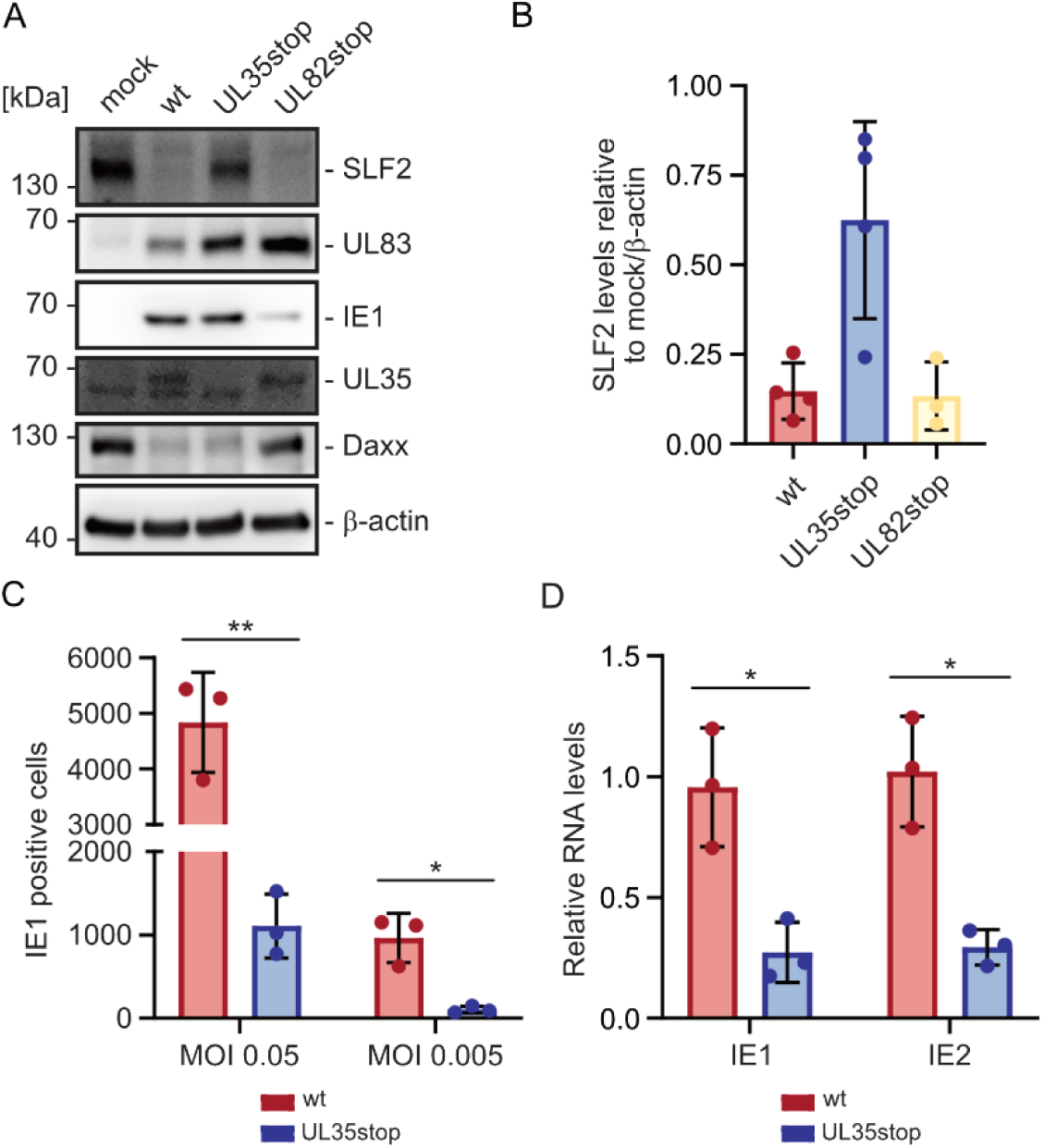
Analysis of SLF2 degradation and initiation of viral gene expression after infection with a recombinant virus defective for expression of UL35. (A-B) UL35-mediated degradation of SLF2 in an infection context. HFF cells were either mock infected or infected for 6 hours with wildtype TB40/E (wt) or a TB40/E virus with a stop codon in UL35 (UL35stop) which abrogates UL35 expression but leaves UL35a expression intact (MOI of 1). Viral inocula were titrated for equivalent IE1 expression. TB40/E UL82stop (UL82stop) was included as additional control. (A) Western blot analysis to detect the levels of SLF2 in whole cell lysates. The viral proteins UL83, IE1 and UL35 and the cellular protein Daxx were included as controls; β-actin was analysed as a loading control. (B) The relative protein levels of SLF2 in infected cells were quantified and are displayed as fold of mock normalized to β-actin. Data are shown as mean values +/- SD from 3 or 4 biological replicates. (C) The absence of UL35 is associated with a reduction in HCMV infectivity. HFF cells were infected either at an MOI of 0.05 (C) or 0.005 (D) with wildtype TB40/E (wt) or TB40/E UL35stop (UL35stop) using viral inocula carefully titrated for an equivalent nuclear uptake of viral genomes. After 24 hours, the numbers of IE1 positive cells were counted using the ImageXpress Pico Automated Cell Imaging System (Molecular Devices) and the means are reported +/- the standard error of the mean (SEM) from three biological replicates. Statistical analysis was performed using Welch’s two-sided unpaired t-test ** p<0.01, * p<0.05. (D) The absence of UL35 is associated with a defect in immediate early transcription. Total cellular RNA was extracted 8 hours post infection with either wildtype TB40/E (wt) or TB40/E UL35stop (UL35stop) (MOI 0.1) using viral inocula titrated for an equivalent nuclear uptake of viral genomes. RT-qPCR was used to determine mRNA levels of IE1 and IE2. Values were normalized to the reference gene GAPDH and are shown as mean values from three independent biological replicates +/-SD. Statistical analysis was performed with Welch’s two-sided unpaired t test, * p<0.05.

### Increased association of vDNA with SMC5/6 containing PML-NBs in the presence of SLF2

To obtain further evidence for a direct repression of viral transcription we decided to study the association of incoming viral genomes with SMC5/6. Previous studies demonstrated that entrapment of viral DNA (vDNA) by PML-NBs can serve as a proxy for genome silencing [17]. We first investigated the colocalization between the SMC5/6 complex and PML-NBs after infection with TB40/E UL35stop. At 6 hpi, wildtype TB40/E and TB40/E UL82stop infection resulted in a complete dissociation of SMC5/6 from PML-NBs (Fig. 8 A, B). In contrast, SMC5/6 localization at PML-NBs was preserved after infection with TB40/E UL35stop (Fig. 8A, B). Of note, both the abrogation of UL35 and pp71 (UL82) expression resulted in morphological changes of PML-NBs including enlargement and the occurrence of ring-like structures (see Fig. 8A, PML). Strikingly, however, the lack of UL35 correlated with a significant increase in the number of enlarged SMC5/6 foci (> 0.5 µm in diameter) at PML-NBs considerably exceeding the association between SMC5/6 and PML-NBs in mock cells (Fig. 8 A, C). This suggests that viral infection may trigger the localization of SMC5/6 at PML-NBs and this is antagonized by UL35-instituted SLF2 degradation.

**Fig. 8:**
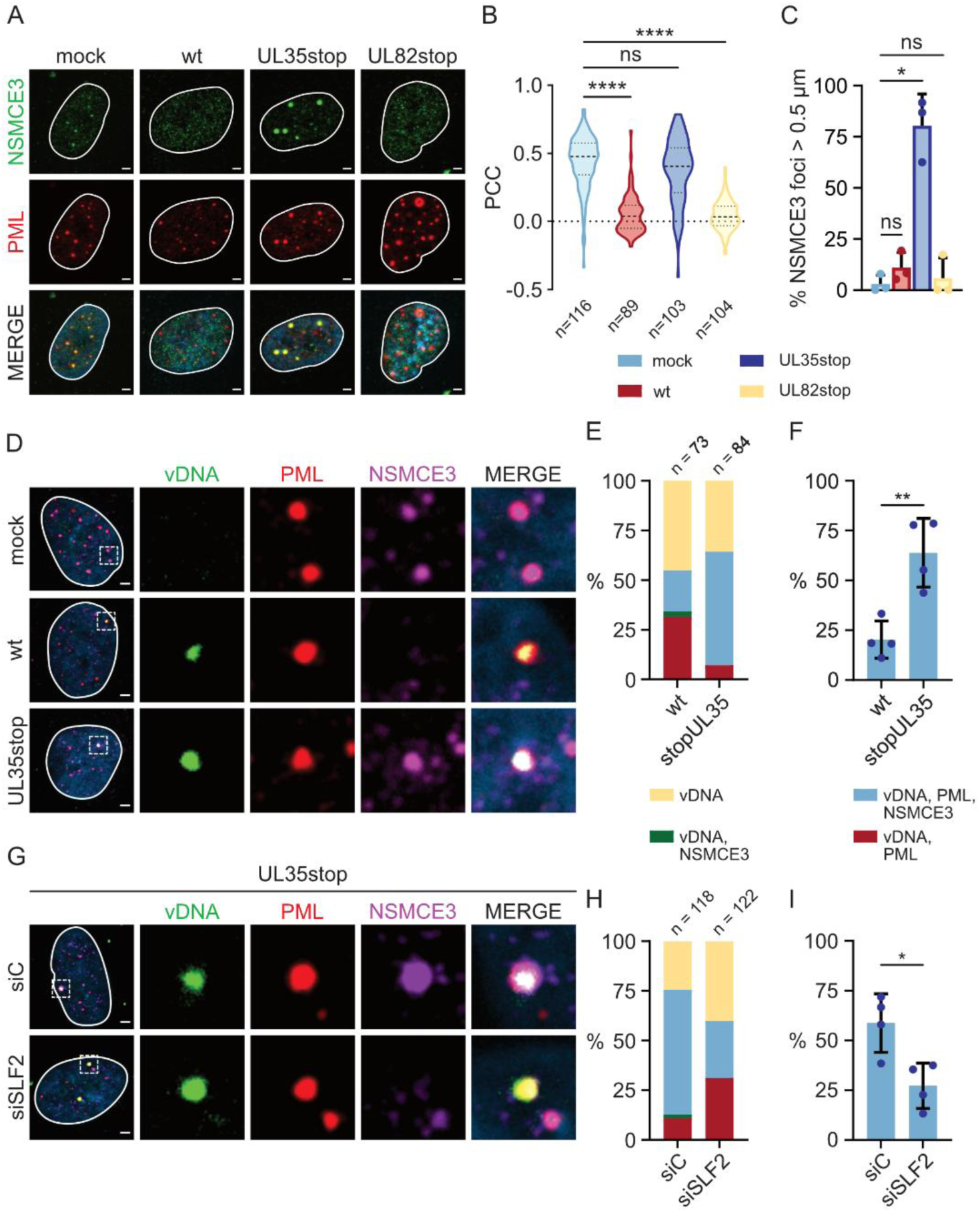
SMC5/6 associates with parental viral genomes in a SLF2-dependent manner. (A-C) Colocalization of SMC5/6 and PML after infection with HCMV defective for UL35 expression. HFF cells were infected (MOI 1) with TB40/E wt (wt), TB40/E UL35stop (UL35stop), TB40/E UL82stop (UL82stop) or were left uninfected (mock). (A) Indirect immunofluorescence analysis of cells harvested at 6 hpi to investigate the colocalization between NSMCE3 and PML. (B) Colocalization analysis of PML and NSMCE3 via quantification of PCCs. The violin plot summarizes the results derived from three independent biological replicates. The median, the first and the third quartile are indicated. (C) Accumulation of enlarged NSMCE3 foci after infection with TB40/E UL35stop. ImageJ-based quantification of NSMCE3 foci with a diameter > 0.5 µm was performed using images of at least 130 cells derived from three independent experiments. Foci > 0.5 µm were grouped and reported as percent of the total number of counted foci. Statistical test: one-way ANOVA (Kruskal-Wallis test). ns: non-significant, * p<0.05, **** p<0.0001. (D-F) Increased association of SMC5/6 with viral DNA (vDNA) in the absence of SLF2 degradation. HFFs were left uninfected (mock) or were infected with either EdC-labeled TB40/E wt or TB40/E UL35stop (MOI 0.01, titrated for viral genome equivalents). After 4 hours, cells were subjected to immunofluorescence staining of PML and NSMCE3, followed by click chemistry to visualize EdC-labeled vDNA. (D) Colocalization of vDNA, PML and NSMCE3. A representative image is shown together with a magnification of the vDNA-inset in the different channels. Scale bar: 2 µm. (E) A total of more than 70 vDNA foci were analysed and grouped into the categories vDNA alone (vDNA), colocalization with NSMCE3 (vDNA, NSMCE3), colocalization with PML (vDNA, PML) or colocalization with NSMCE3 and PML (vDNA, PML, NSMCE3). The graph depicts the percentage of each vDNA foci category. (F) Increase of colocalization of vDNA foci with both NSMCE3 and PML after infection with TB40/E UL35stop. The bar graph shows the means +/-SD of four biological replicates. Statistical analysis was performed using Welch’s t two tailed t test. **, p < 0.01. (G-I) SLF2 depletion reverses the increased association of vDNA with NSMCE3 after infection with TB40/E UL35stop. HFFs were transfected either with control siRNA (siC) or an siRNA directed against SLF2 (siSLF2) followed by infection with EdC-labeled TB40/E UL35stop. The colocalization of PML, NSMCE3 and vDNA was analyzed as described for panel D. (G) A representative image is shown together with a magnification of the vDNA-inset in the different channels. Scale bar: 2 µm. (H) Colocalization categories of the detected vDNA foci as described for panel E. More than 110 vDNA foci were analysed and the graph depicts the relative percentage of each category. (I) SLF2 depletion decreases the colocalization of vDNA foci with both NSMCE3 and PML after infection with TB40/E UL35stop. The bar graph shows the means +/- SD of four biological replicates. Statistical analysis was performed using Welch’s t two tailed t test. *, p < 0.05.

To study the entrapment of vDNA by SMC5/6 containing PML-NBs, genomes of TB40/E wildtype and UL35stop viruses were labeled with EdC as described previously [17]. At 4 hpi, HFF cells were stained for PML and NSCME3, serving as marker proteins for PML-NBs and SMC5/6 complexes, respectively. Click chemistry was then used to visualize vDNA followed by a colocalization analysis of both protein complexes with genomes (Fig. 8D-F). In addition to an overall increment of vDNA at PML-NBs, we observed a significant increase in the number of vDNA foci associated with SMC5/6 containing PML-NBs after infection with TB40/E UL35stop (Fig. 8F). To demonstrate that this depends on SLF2, cells were transfected with either a control siRNA (siC) or a SLF2 specific siRNA (siSLF2) followed by infection with TB40/E UL35stop. Indeed, we observed that siRNA-mediated depletion of SLF2 reversed the effect of UL35 on the colocalization of vDNA with SMC5/6 and PML-NBs (Fig. 8G-I). We observed an overall decrease in PML-NB association. In particular, the number of SMC5/6 containing PML-NBs at vDNA was significantly reduced (Fig. 8I). In conclusion, these findings provide further evidence for a host-directed mechanism of viral genome repression that is antagonized by UL35.

## Discussion

In this study we identify the HCMV tegument protein UL35 as a novel antagonist of SMC5/6-mediated gene silencing. The SMC5/6 complex has multiple important functions in maintaining the integrity of the cellular genome such as its contribution to various DNA repair processes [34]. Beyond this, accumulating evidence demonstrates that DNA manipulation by SMC5/6 selectively represses the transcription of circular but not of linear DNA [4]. This was first described with hepatitis B virus which persists as an extrachromosomal circular DNA in infected cells [35]. Further studies revealed that not only viral circular DNA, but also plasmids can serve as substrates for SMC5/6-mediated repression [4]. As a physical basis for this, a preferential binding of SMC5/6 to circular rather than linear extrachromosomal DNA was detected [36]. Notably, superhelical stress generated by transcription appears to serve as an important stimulus for SMC5/6 association and subsequent DNA compaction [36, 37]. This represents an elegant mechanism of host defense against circular DNA viruses triggered by active gene expression [36].

Thus, it is not surprising that viruses developed mechanisms to evade this host defense in the evolutionary arms race. For most viruses known to antagonize SMC5/6, including HBV, adenoviruses and the herpesviruses EBV and KSHV, components of the core complex were identified as targets for degradation [11, 12, 35, 38, 39]. The core structure consists of a SMC5/SMC6 heterodimer and several ‘non-SMC’ proteins or ‘elements’ (NSMCE) that form a stable holo-complex (Fig. 1A) [7]. Upon investigation of SMC5/6 protein abundance after HCMV infection, we could not detect a downregulation of SMC5/6 core proteins (Fig. 1B). Instead, we observed decreased expression of the SMC5/6 associated protein SLF2 (Fig. 1B). SLF2 was first described as component of a subcomplex comprising SLF1, SLF2 and RAD18 which recruits SMC5/6 to DNA lesions to suppress genome instability [40]. Consistent with a critical function of SLF2 for genome integrity, mutations in SLF2 cause a neurodevelopmental disease named Atelis syndrome which is characterized by microcephaly, short stature and chromosomal breakage [41]. Importantly, research on SMC5/6 repression of HBV gene expression revealed a second function of SLF2 linking SMC5/6 to a different multiprotein complex implicated in intrinsic antiviral immunity, the subnuclear structure PML-NBs [42]. First evidence for this was obtained in a study demonstrating a colocalization of the SMC5/6 complex with PML-NBs in primary human hepatocytes and depletion of PML-NB proteins induced HBV gene expression suggesting an interconnection between SMC5/6 and PML-NB instituted silencing [43]. Subsequently, SLF2 was identified as the responsible factor which interacts with SMC5/6 and directs HBV cccDNA to PML-NBs for entrapment and transcriptional repression [26]. Consistently, we observed a colocalization of SMC5/6 and SLF2 with PML-NBs in primary human fibroblasts (Fig. 2, Fig. 6E). Both SLF2 depletion and HCMV infection abrogated this colocalization (Fig. 2). Furthermore, in the absence of viral antagonization, enhanced entrapment of parental HCMV DNA in SMC5/6 containing PML-NBs was detected correlating with a defect in the onset of immediate-early gene expression (Fig. 7 and 8). Interestingly, we observed enlarged SMC5/6 accumulations at PML-NBs after infection with an antagonization-defective HCMV suggesting a virus-triggered mechanism of recruitment (Fig. 8C). Since depletion of SLF2 reversed the enhanced entrapment of HCMV DNA in SMC5/6 containing PML-NBs (Fig. 8G-I), this suggests a conserved role of SLF2 in viral genome silencing. Moreover, a recent study proposed the regulation of SMC5/6 by two mutually exclusive SLF2-containing subcomplexes – SLF1/2, directing SMC5/6 to chromosomal DNA lesions, and SIMC1-SLF2, recruiting SMC5/6 for repression to PML-NBs which appears to be antagonized by the SV40 large T antigen [9]. However, in contrast to HCMV, which induced a downregulation of SLF2 via proteasomal degradation (Fig. 3E-G), neither HBV nor SV40 affected the expression levels of SLF2. So far, depletion of SLF2 has only been described in the context of HIV-1 infection increasing the chromatin accessibility of unintegrated lentiviral DNA [44]. Thus, HCMV appears to be the first bona fide DNA virus which evolved SLF2 degradation as a mechanism to antagonize SMC5/6 instituted silencing.

Many viral effector proteins that modulate SMC5/6 have previously been identified as enigmatic promiscuous transactivators. For instance, the HBV protein HBx was reported to activate viral enhancers as well as various RNA polymerase II and III promoters in the experimental setting of transient expression analysis [45, 46]. Later on, it was recognized that only extrachromosomal DNA templates were affected which is now well explained by the finding that HBx targets SMC5/6 for degradation to induce a general relieve of plasmid repression [35, 47]. Using a targeted screening approach, we identified the tegument protein UL35 of HCMV as the viral effector protein inducing SLF2 degradation (Fig. 5). Initial studies on UL35 detected a transactivation capacity of this protein in transient expression assays that was augmented by co-expression of the tegument protein pp71 (UL82) [48]. Furthermore, both proteins were found to co-localize at PML-NBs [48]. The tegument protein pp71 (UL82) is meanwhile well recognized for its capacity to antagonize PML-NB instituted repression of viral transcription via disruption and proteasomal degradation of the ATRX/Daxx histone H3.3 chaperone complex [18, 49–51]. Since pp71 together with UL35 were described to direct the proteasomal degradation of yet another restriction factor, the cellular BclAF1 protein, we speculated that pp71 might also be important for SLF2 degradation [52]. However, infection experiments with a recombinant virus harboring a stop codon in UL82 resulted in degradation of SLF2 while Daxx levels were sustained (Fig. 4A, 7A). Vice versa, abrogation of UL35 expression preserved SLF2 expression while downregulating Daxx protein levels, clearly demonstrating that HCMV has evolved two different effector proteins to target either SLF2 or Daxx separately for degradation (Fig. 7A). However, although no SLF2 downregulation was found upon transient co-expression with pp71 (UL82), we noticed a colocalization of both proteins in ring-like subnuclear structures (Fig. 5A, B). It needs to be investigated in further studies whether this indicates a functional interrelation between both proteins. Interestingly, the EBV effector protein BNRF1 also attacks the SMC5/6 complex via Cullin-7 dependent degradation of SMC5/6 core components, and at the same time disrupts the histone H3.3 chaperone complex Daxx/ATRX to antagonize host-cell intrinsic defenses [11, 53]. This indicates a convergent evolution of herpesviral effector proteins to target both SMC5/6 and PML-NBs.

Of note, proteomic profiling revealed an interaction of UL35 with three components of the CRL4-DCAF1 E3 ubiquitin ligase complex, DCAF1, DDB1 and DDA1, previously shown to be targeted by the HIV-1 Vpr protein [54]. Furthermore, both Vpr and UL35 induce a DNA damage response resulting in the accumulation of cells in the G2 phase of the cell cycle [54, 55]. Since Vpr was identified as the HIV-1 encoded effector protein responsible for SLF2 degradation, which we confirmed in our study (Fig. 6A and B), we assume that both UL35 and Vpr hijack the CRL4-DCAF1 E3 ligase to induce the ubiquitination and subsequent degradation of SLF2 [44]. It remains to be determined whether interaction with CRL4-DCAF1 is also relevant for the reported activities of UL35 to degrade the restriction factor BclAF1 or the basement protein nidogen 1 implicated in neurological damage induction by congenital CMV infection [52, 56, 57].

However, protein degradation is not the only mode of action of the multifunctional UL35 effector protein. For instance, UL35 has been shown to antagonize type I interferon signaling via interacting with TANK-binding kinase 1 and this was not dependent on CRL4-DCAF1 [28]. Furthermore, it is well known that the UL35 gene region encodes two proteins, named UL35 and UL35a, which distinct functions [29]. The 75-kDa UL35 protein is packaged into the tegument of virions and thus imported into cells. In contrast, the smaller 22-kDa UL35a is expressed predominantly late during infection. It is not contained in viral particles but plays an important role for the assembly of mature particles via translocating the tegument proteins pp71 (UL82) and pp65 (UL83) from the nucleus to the cytoplasm [31–33]. This impedes the phenotypical characterization of HCMV mutants exhibiting a deletion of the entire UL35 gene region due to a severe replication defect caused by impaired virus assembly [32, 33]. Since our results indicated that UL35a lacks the capacity to degrade SLF2, we utilized an HCMV mutant harboring stop codons within its reading frame which abrogate UL35 but not UL35a expression (Fig. 6 and 7). Consistent with a previous publication, we observed reduced immediate-early gene expression upon infection under conditions of standardized viral DNA (Fig. 7C and D) [32]. Together with our data on enhanced entrapment of viral progeny DNA which is reversed upon SLF2 depletion (Fig.8), this demonstrates that UL35, but not UL35a, serves as a novel antagonist of SMC5/6 mediated gene silencing. Since SLF2 exerts dual functions during gene silencing and DNA repair, depletion of this important cellular protein by an HCMV tegument protein may also be relevant for the reported induction of chromosomal defects by HCMV infection [56, 58].

## Material and methods

### Antibodies, expression plasmids, primers and compounds

Key resources used in this study are listed in S1 Table and S2 Table. Oligonucleotide primers were purchased from Biomers GmbH (Ulm, Germany). SiRNAs were obtained from Horizon Discovery (Cambridge, UK).

### Plasmid constructions

All primers used for cloning purposes are listed S2 Table. To generate plasmids expressing either HA-tagged pp71 (UL82-HA) or pp65 (UL83-HA) the respective coding sequences were amplified by PCR with primer pairs HA_pp71_EcoRI-fw/pp71_XhoI-rev and UL83_BamHI-fw/UL83(561)_HA-XhoI-rev, respectively, followed by restriction enzyme digestion and insertion into vector pcDNA3.0 (NovoPro BioScience, Shanghai, China). Expression plasmids for FLAG-tagged UL35 and UL35a were also constructed via PCR with primer pairs UL35-ClaI-fw/UL35-XhoI-rev and UL35A-ClaI-fw/UL35XhoI-rev and subsequent insertion into vector pHM971 [59]. Vectors for dox-inducible expression of FLAG-tagged UL35 and UL35a were generated by PCR amplification of the respective coding sequences with primer pairs FLAG_UL35_NotI-fw/UL35_MluI-rev and FLAG_UL35a_BamHI-fw(45-859/UL35_NotI-rev and insertion into pLVX-Tight-Puro (Clontech).

### Cells and viruses

All cells were cultured at 37°C and 5% CO_2_. Primary human foreskin fibroblasts (HFFs), isolated from human foreskin tissue as described previously, and human retinal pigment epithelial cells ARPE-19 (ATCC-CRL-2302) were cultivated in Dulbeccós modified eagle medium (DMEM) (Gibco) supplemented with 5% fetal bovine serum (FBS) (Capricorn) and penicillin-streptomycin (Sigma-Aldrich) [60]. Human embryonic kidney HEK293T cells (DSMZ ACC 635) and doxycycline (dox)-inducible UL35-HA together with control cells were cultured as described previously [17, 28]. Induction of protein expression was achieved by the addition of dox (Sigma) for 24 hours (h) at concentrations of 0.5 -1 µg/ml. Cells were tested for the absence of mycoplasma contamination using the MycoAlert Mycoplasma Detection Kit (Lonza) as described by the manufacturer.

All viruses used in this study are listed in S1 Table. The recombinant HCMV TB40/E UL82stop was constructed by markerless mutagenesis with primers kindly provided by C. Sinzger (Ulm, Germany) [61, 62]. Successful recombination was confirmed by restriction enzyme digestion, PCR and nucleotide sequence analysis of the recombined gene region. Infection experiments of HFF cells were performed either with wildtype or recombinant HCMVs based on strain TB40/E as described previously [17, 63]. For ARPE-19, centrifugation of the viral inoculum for 30 min at 1200 x g was performed to enhance infection efficacy. UV inactivation of viral inocula was performed by UV irradiation (0.12 J/cm^2^ for 3 min) using a CL-1000 Ultraviolet Crosslinker. For treatment of infected cells with the inhibitors, the compounds (listed in S1 Table) were added immediately upon infection and at medium change. The following inhibitors were used: cycloheximide (150 µg/ml), MG132 (10 µM), MLN4924 (10 µM), PYR-41 (5 µM) (see also S1 Table).

For titration of HCMV, HFFs were infected with serial dilutions of the virus in 96 well plates. 24 hours post infection (hpi), cells were fixed with 80% acetone and stained for IE1. Automated quantification of IE1-positive cells was performed using the ImageXpress Pico System (Molecular Devices), and viral titers were calculated in immediate-early (IE) units/ml. Titration based on the quantification of intracellular viral genomes was performed with a Taqman real-time PCR as described in [64]. Briefly, HFF cells were seeded in triplicates in 6well plates and infected at an MOI of 0.1. After 16 h, genomic DNA was extracted using the DNeasy Blood & Tissue Kit (Qiagen) followed by the quantification of viral genomes and cellular albumin by real-time PCR exactly as described previously [63]. For the determination of viral genome equivalents, the ratio of the IE1 and albumin was calculated and averaged for each triplicate. These calculations were used to normalize infection conditions to the delivery of equivalent intracellular genome copy numbers.

### Transfection

HFF or ARPE-19 cells were seeded at a density of 1 × 10^5^ cells per well of a 12 well plate in 0.8 ml DMEM supplemented with 5% FBS. On the next day, cells were incubated for 1 h with 8 µl K2 multiplier (Biontex) followed by addition of plasmid DNA (1 µg/well) with 3 µl of Lipofectamine 2000 (ThermoFisher Scientific), diluted in Opti-MEM (Gibco) according to the manufacturers protocol. For siRNA transfection, a final concentration of 15 nM of siRNA and 6 µl of Lipofectamine RNAiMAX (ThermoFisher Scientific) in OptiMEM was added to each well. For all transfections, the liposome solution was added dropwise to cells, removed after 5 h incubation and replaced by DMEM supplemented with 5% FBS and penicillin-streptomycin. Plasmid DNAs and siRNAs used in this study are listed in S1 Table and S2 Table, respectively.

### Lentiviral transduction

Primary HFF expressing Dox-inducible UL35-HA or the empty vector control (ev) were generated by lentiviral transduction using pW-YC1hygro expression constructs (Tet-On system), the packaging plasmid psPAX2 and the envelope plasmid pMD2.G. Plasmids psPAX2 and the envelope plasmid pMD2.G were gifts from D. Trono (Addgene #12260 and #12259. The blasticidin-resistance cassette in the pW-Y1blast vector (kindly provided by L. Dölken and T. Hennig, Hannover Medical School) was replaced with a hygromycin resistance cassette to generate pW-YC1hygro [65]. Transduced cells were selected in the presence of 50 µg/ml hygromycin B (Invitrogen). Dox-inducible HFFs for the expression of UL35-FLAG and UL35a-FLAG cells were generated with the Lenti-X™ Tet-On® Advanced Inducible Expression System (Clontech) according to instructions of the manufacturer. Transduced cells were selected in the presence of 500 µg/ml geneticin (Invivogen) and 5 µg/ml puromycin (Invivogen). MYC-SLF2 overexpressing cells were produced essentially as described in [66]. Briefly, replication-deficient lentiviral particles were generated by transfection of HEK293T cells with the lentiviral expression plasmid pLVX-Neo 1xmyc wt slf2 siRNA-res (kindly provided by G. S. Stewart, Birmingham, UK) together with packaging plasmids pCMVdeltaR8.9 and pLP-VSVg (listed in S1 Table) [41]. The day after transfection, fresh medium was provided. After two days, the lentiviral-containing supernatant was cleaned from cell debris by filtration through a 0.45 µm sterile filter and used to directly transduce target cells, or stored at -80°C. To transduce HFF cells, 8 × 10^4^ cells/well were seeded in 6-well plates. One day later, the cells were incubated with lentivirus supernatant in the presence of 7.5 µg/ml polybrene (Sigma-Aldrich) for 24 h. After addition of fresh cell culture medium, the selection of successfully transduced cells was initiated by the addition of 500 µg/ml geneticin.

### Western blotting

Western blot analysis was performed essentially as described [64]. Briefly, cells were harvested and resuspended in PBS and 4 x sodium dodecyl sulfate-polyacrylamide gel electrophoresis (SDS-PAGE) loading buffer followed by boiling for 10 min at 95°C and sonication for 1 minute. Proteins were separated on SDS gels containing 6 to 15% polyacrylamide before being transferred to PVDF membranes (BioRad). After incubation with the target antibodies listed in S1 Table, the proteins were visualized through chemiluminescence reaction using a FUSION FX7 imaging system (Vilber). Signal intensities were quantified with ImageLab software (BioRad) and expressed as fold of the chosen reference.

### Immunofluorescence

The indirect immunofluorescence protocol was adapted from [20]. Briefly, coverslips (VWR) were coated for 30 min at 37°C and 5% CO_2_ with 0.1% gelatine (Sigma) in H_2_O prior to seeding of the cells. Cells were fixed with 4% paraformaldehyde (PFA) (Sigma-Aldrich) for 10 min at room temperature at the designated time point and quenched by incubating the cells with a solution of 50 mM Glycine (Sigma-Aldrich) and 50 mM NH_4_Cl (Merck) in PBS. The permeabilization was performed with 0.1% Triton X-100 (Sigma) in PBS at 4°C for 15 min, followed by blocking with 2 mg/ml gamma-globulin from human blood (Sigma-Aldrich) for 30 minutes at 37°C. Target proteins were stained with the primary antibodies diluted in PBS supplemented with 1% FBS for at least 1 h at 37°C. Alternatively, incubation was performed for 24 h at 4°C. After extensive washing the samples were incubated with secondary antibodies for 30 min at 37 °C. Finally, the samples were washed with PBS and mounted with Vectashield DAPI Mounting Medium (Vector Laboratories) on microscope slides (Epredia). All antibodies used in this study are listed in S1 Table. Images were acquired with a Zeiss Axio Observer Z1 with an Apotome.2 and analysed with the ZEN 3.0 (Blue edition) software (Carl Zeiss Microscopy). The antibodies used are listed in S1 Table.

### Viral genome labelling and click chemistry

Immunofluorescence-based visualization of viral genomes was performed similarly as described in [17]. Briefly, HCMV strain TB40/E was propagated in HFFs until the appearance of a distinct cytopathic effect. Two days before harvesting, the medium was replaced every 24 h with new medium supplemented with 0.5 µM 5-ethynyl-2′-deoxycytidine (EdC) (Sigma). Afterwards and every second day until complete cell lysis, the viral supernatant was collected and new medium with 0.5 µM EdC was added. The harvested supernatants were centrifuged for 10 min at 2800 rpm to remove cellular debris. Viral particles were concentrated by ultra-centrifugation at 23.000 rpm for 70 min at 10° C.

Afterwards, the viral pellet was washed with 4 ml of fresh medium without EdC followed by gentle shaking for 2 h at 4°C. Finally, the pellet was resuspended with a 20 gauge syringe needle (Sterican), aliquoted and stored at -80 °C. The labelling efficacy was assessed through immunofluorescent staining of the viral tegument protein UL32, serving as a marker for viral particles, at 4 hpi of HFFs. To visualize the EdC-containing viral DNA, a click-chemistry based reaction was performed as described [17]. Briefly, after incubation with the secondary antibody, the samples were washed with PBS and incubated for 2 h at room temperature in the dark with a freshly prepared click chemistry solution composed of 0.01 M PBS, 10 µM AlexaFluor Azide 488 (ThermoFisher), 10 mM (+)-sodium L-ascorbate (Sigma), 1 mM copper(II) sulfate pentahydrate (CuSO_4_) (Sigma), 10 mM aminoguanidine hydrochloride (Sigma-Aldrich) and 1 mM Tris-hydroxypropyltriazolylmethylamine (THPTA) (Sigma-Aldrich). Afterwards, the samples were washed with PBS, incubated for 5 min with 3% BSA (Sigma-Aldrich) solution at room temperature and finally washed with PBS before mounting the coverslips on microscope slides using Vectashield DAPI Mounting Medium (Vector Laboratories). The labelling efficiency was calculated as the number of pUL32 signals colocalizing with EdC signals.

### Colocalization analysis and focus quantification

Calculation of the Pearson’s correlation coefficient (PCC) was performed with the colocalization module of ZEN blue v2.3 (Carl Zeiss Microscopy). In brief, a rectangular area delimiting each single cell nucleus was selected with the drawing tool and the pictures were analysed with ZEN 2.3 (Carl Zeiss Microscopy). The thresholds for the quantification were defined in a way to encircle the areas of interest using the relative signal intensities as excluding criteria. Immunofluorescence quantification of nuclear foci dimension was performed with Fiji [67]. Briefly, through the function “Analyze Particles” nuclei coordinates are saved. Afterwards, the foci of interest were identified for each nucleus through “Analyze Particles” in the respective picture of the target immunofluorescent channel. Subsequently, the area and mean fluorescence intensity of the identified foci were measured, respectively.

### RNA isolation and reverse transcription PCR (RT-qPCR)

To extract total cellular RNA, 3 × 10^5^ HFF were seeded in 6 wells plate in triplicates. 8 h after infection, cells were lysed in 0.7 ml of TRIzol (Invitrogen) for 5 min at room temperature. Total RNA was isolated using a commercial RNA extraction kit (Zymo Research), and 0.5 µg of the RNA was reverse transcribed using the Maxima FirstStrand cDNA Synthesis Kit (ThermoFisher), by including the 30 min DNase treatment step, as described by the manufacturer protocols. 2 µl of the synthesized cDNAs were analysed through SYBR Green PCR with AriaMx real-time PCR system (Agilent) with a reaction mix composed of SsoAdvanced Universal SYBR Green Supermix (BioRad) and 150 nM primers (listed in S2 Table). The reaction was performed with the AriaMx software v1.7 by doing an initial denaturation step (95°C for 30 s), followed by 40 cycles consisting of a denaturation step (95 °C for 15 s), primer annealing and elongation (60°C for 30 s) and finally a dissociation step (95 °C for 30 s, 65 °C 30 s, 95°C 30 s). The Cq were analysed using the 2^−ΔΔCt^ method, using GAPDH (glyceraldehyde-3-phosphate dehydrogenase) as reference gene.

### Statistical analysis and graphic preparation

Statistical analyses were performed with the built-in analysis of the Prism (Graph Prism v10.6.1 (892)) software. The tests included one-way ANOVA (Kruskal-Wallis test), Mann-Whitney t test, and Wech’s t-test. The results were indicated as marginally significant (p-values < 0.05), significant (p-values < 0.01), and highly significant (p-values < 0.001). Schematic representations were generated using the design software Biorender (www.biorender.com).

## Acknowledgement

We would like to thank Romy Schelling (Ulm, Germany) and Kerstin Laib-Sampaio (Ulm, Germany) for helpful advice and experimental support. Furthermore, we are very indebted to Christian Sinzger (Ulm, Germany) and Grant S. Stewart (Birmingham, UK) for providing recombinant HCMV strains and SLF2 expression plasmids, respectively.

## Author Contributions

Data curation: FM, MS, CS

Conceptualization: TS, MS

Formal analysis: FM, MS, CS, FH, CP, CSE, TM

Funding acquisition: TS, MMB

Investigation: FM, MS, CS, FH, CP, CSE, TM, RR, NA-B

Methodology: CS, FH, TM, RR, NA-B

Project administration: TS, MMB Supervision: TS, MS, NAB, MMB

Visualization: FM, TS

Validation: FM, MS

Writing – original draft: FM, TS

Writing – review & editing: FM, CS, FH, RR, NAB, MMB, TS

## SUPPORTING INFORMATION

**S1 Figure:**
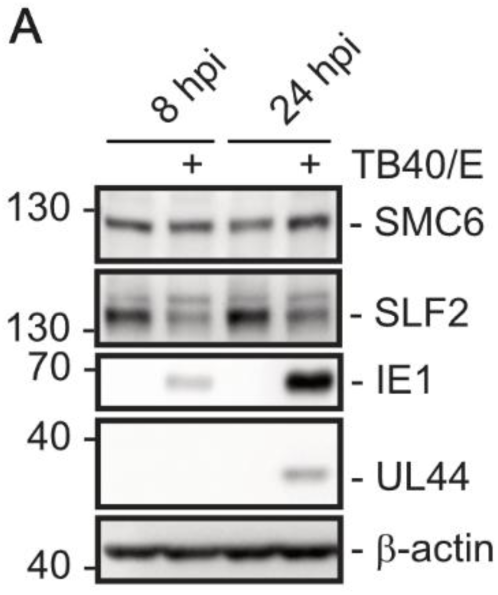
Downregulation of SLF2 upon HCMV infection of ARPE-19 cells. ARPE-19 cells were left uninfected or were infected with HCMV TB40/E (MOI of 1) as indicated. After 8 and 24 h, cells were harvested and SLF2 protein expression was analysed through western blotting. In parallel, expression of SMC6, IE1 and UL44 were analysed; β-actin served as a loading control. Three independent biological replicates were performed.

**S2 Figure:**
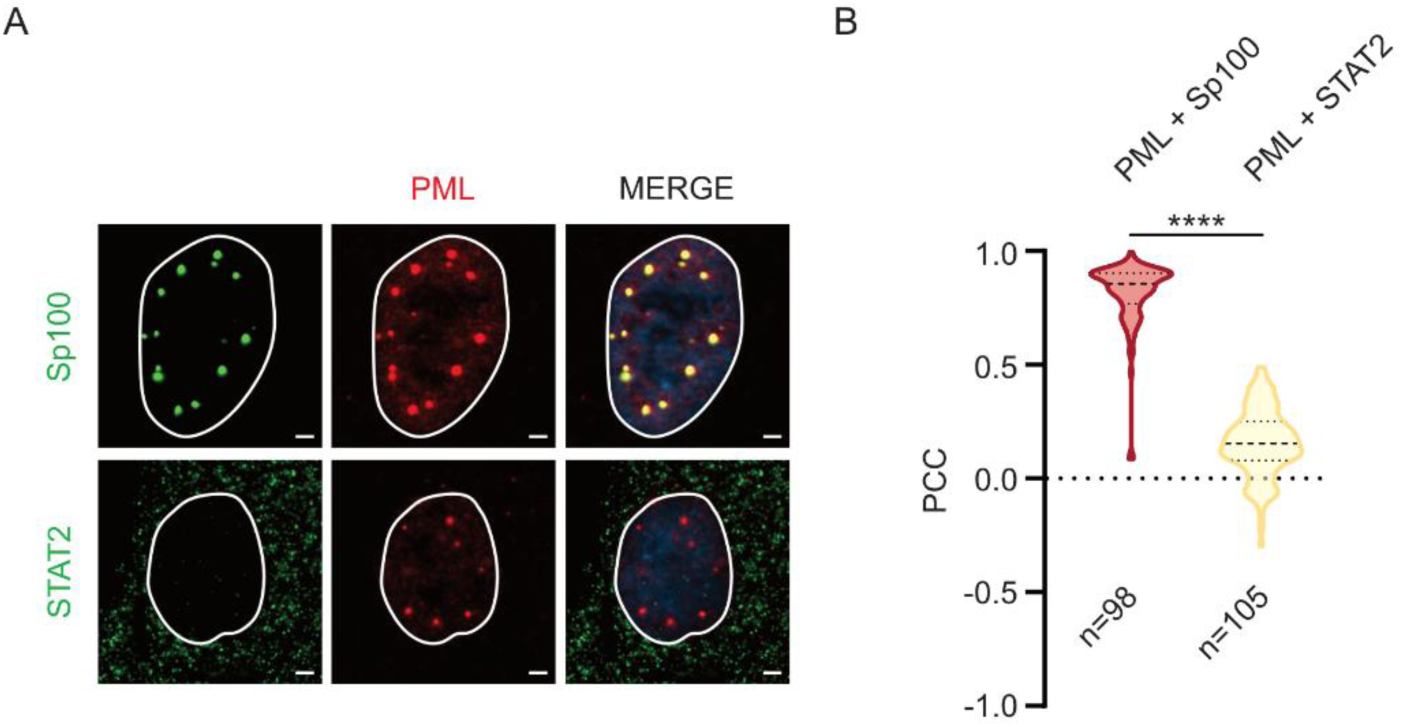
Colocalization analysis via calculation of Pearson’s correlation coefficient (PCC). (A) Immunofluorescence staining of HFF was performed to detect either PML together with Sp100 (used as positive control) or PML together with STAT2 (used as negative control), respectively. (B) The Pearson correlation coefficient (PCC) was calculated with the ZEN Blue colocalization module (Zen 2.3, Carl Zeiss Microscopy GmbH, 2011). To restrict the analysis to the area corresponding to PML-NBs, regions of interest were first drawn around cell nuclei. Afterwards, the maximum white values of the entire picture were identified in the range indicator/histogram panel. The threshold for PML channel and the channel for the putative colocalizing protein was set to 10% and 0% of this value, respectively. A total of more than 90 nuclei were quantified, and the results of three independent biological replicates were grouped in a violin plot. Median, first and third quartile are indicated. The statistical analysis was performed with the Mann Whitney t test, **** p<0.0001.

**S3 Figure:**
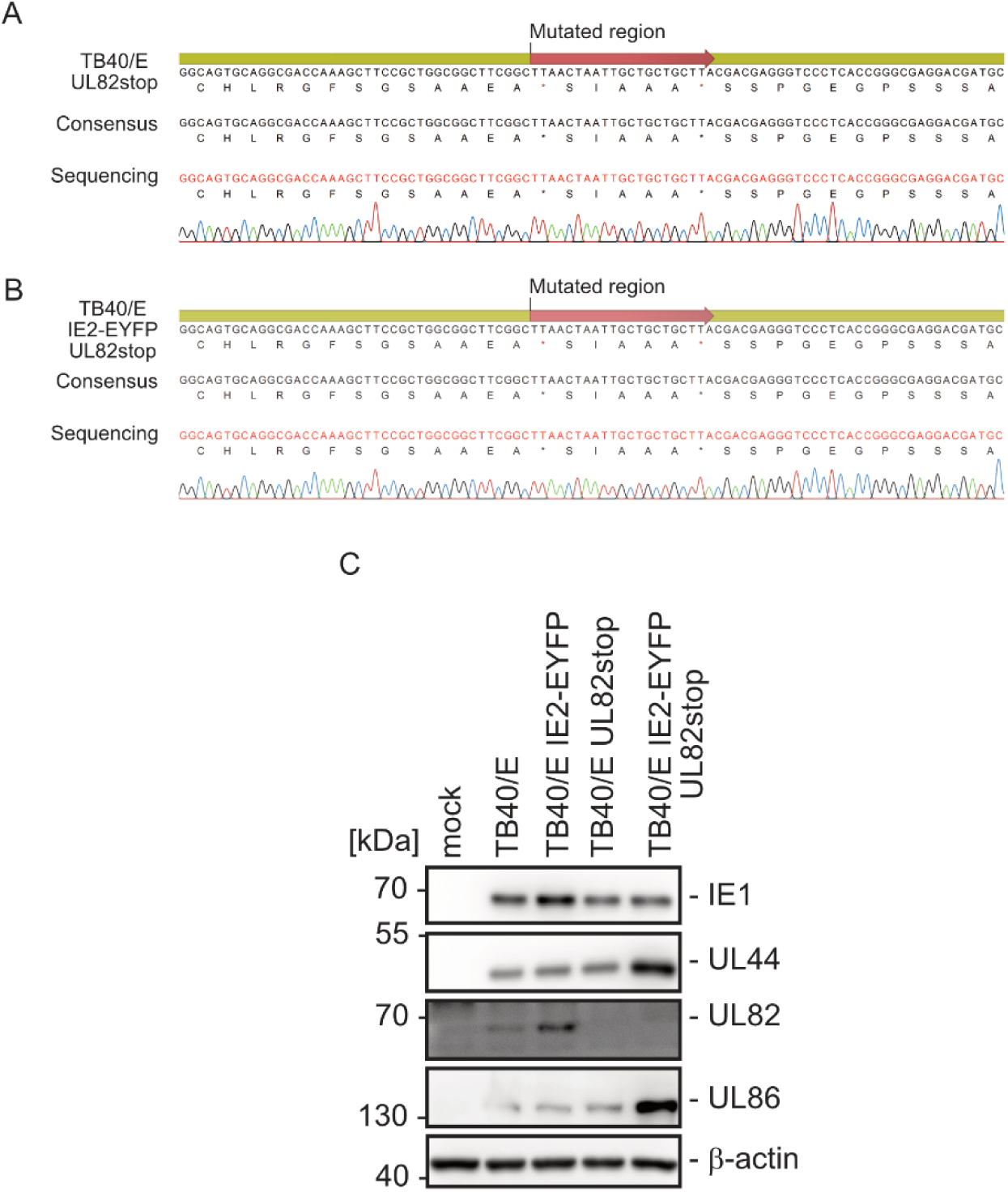
Construction of recombinant HCMVs TB40/E UL82stop and TB40/E IE2-EYFP UL82stop. (A-B) Two stop mutations were inserted into the UL82 gene region of the HCMV BAC TB40/E-BAC4 through markerless *en passant* mutagenesis and were confirmed by nucleotide sequence analysis. The stop codons are indicated by asterisks within the amino acid sequence. The consensus lane shows the results of the pairing between the target sequence and the sequencing results of bacmid DNA containing the mutated viral DNA. (C) Analysis of pp71 (UL82) expression after infection of HFFs with TB40/E wt or the reconstituted TB40/E UL82stop and TB40/E-IE2-EYFP UL82stop (MOI 1). Western blot analysis was performed with cell lysates harvested 5 dpi using a UL82-specific polyclonal antiserum. The immediate early protein IE1, the early protein UL44 and the early-late protein UL86 were detected as controls; β-actin served as a loading control.

**S1 Table:**
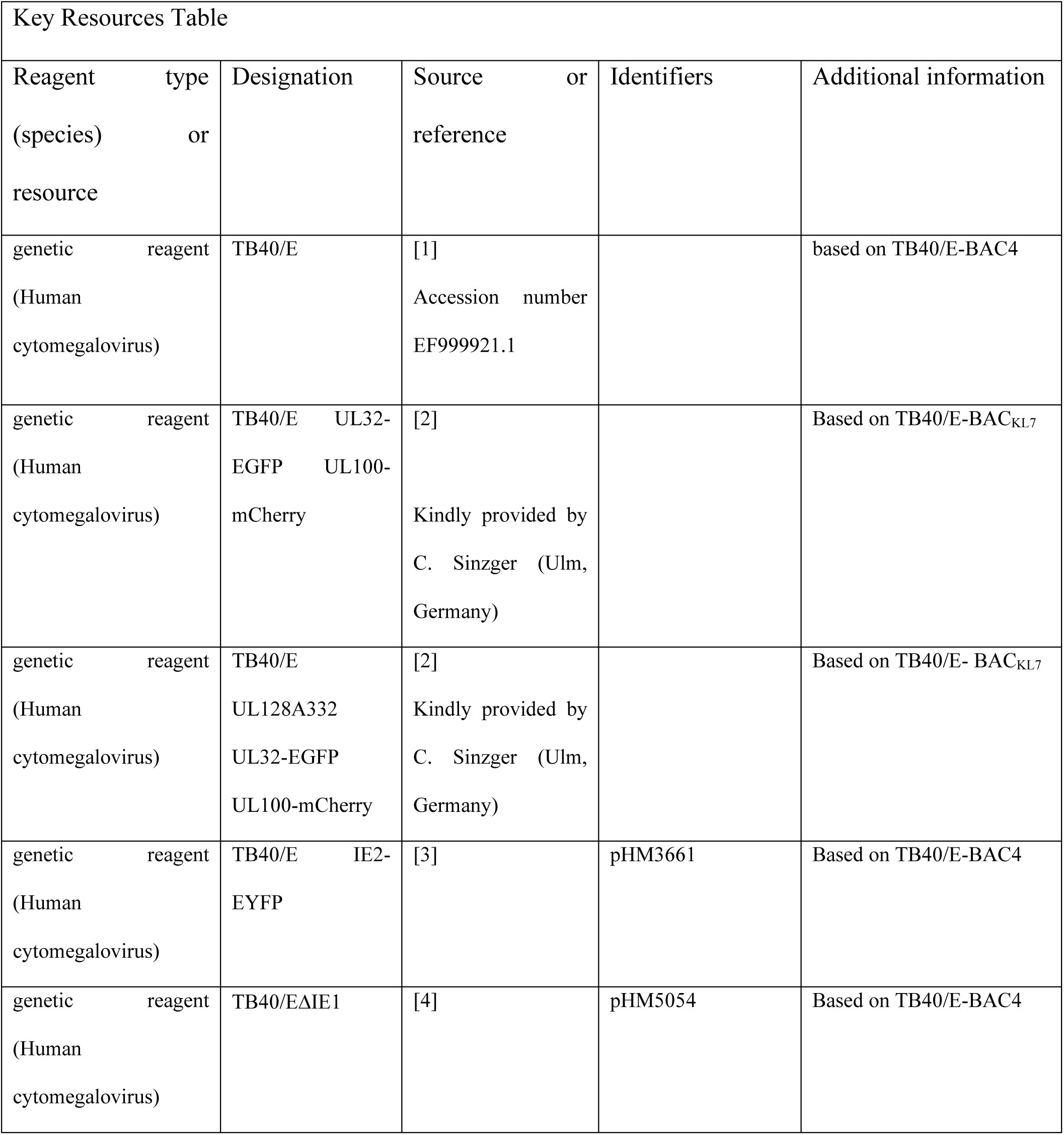
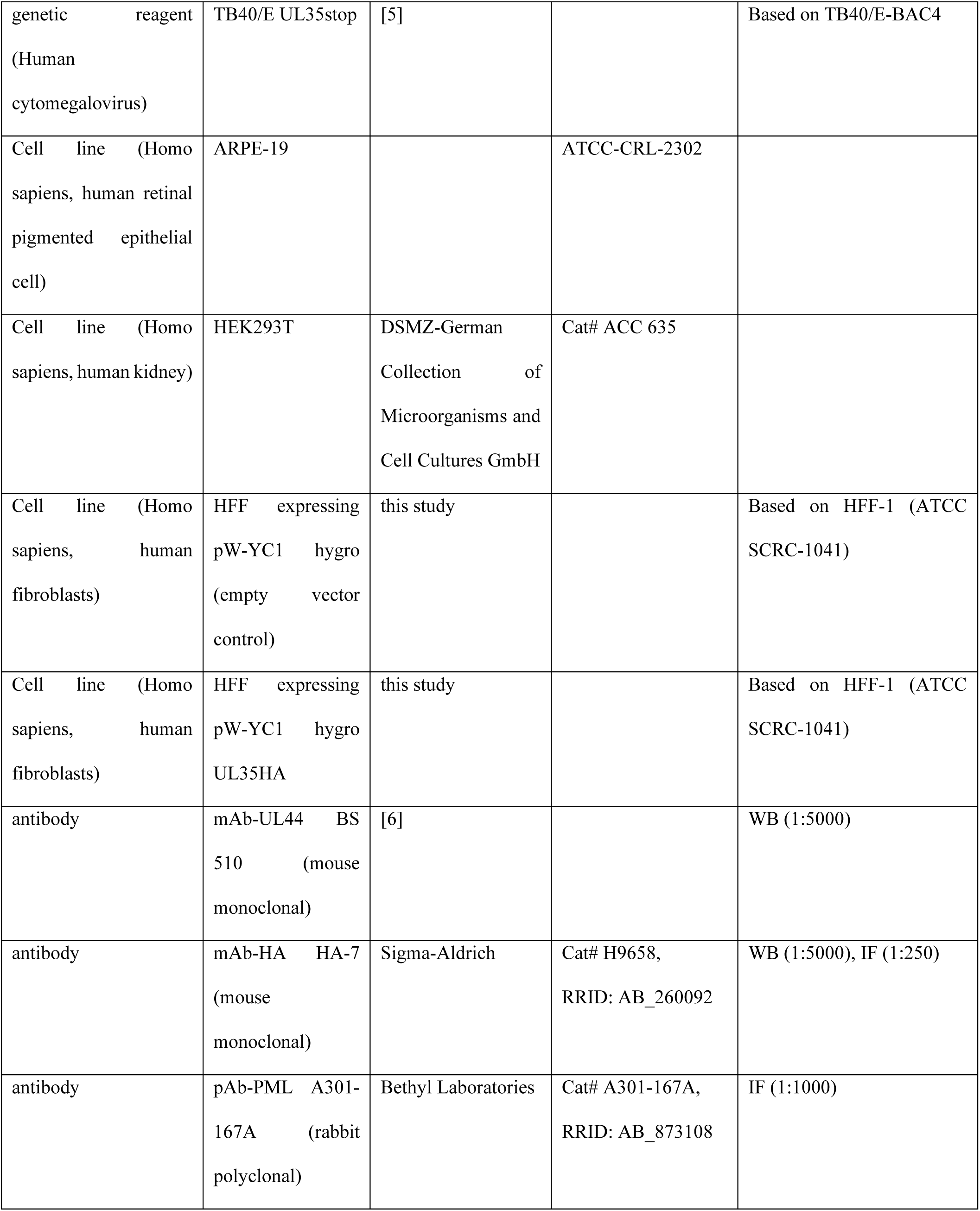
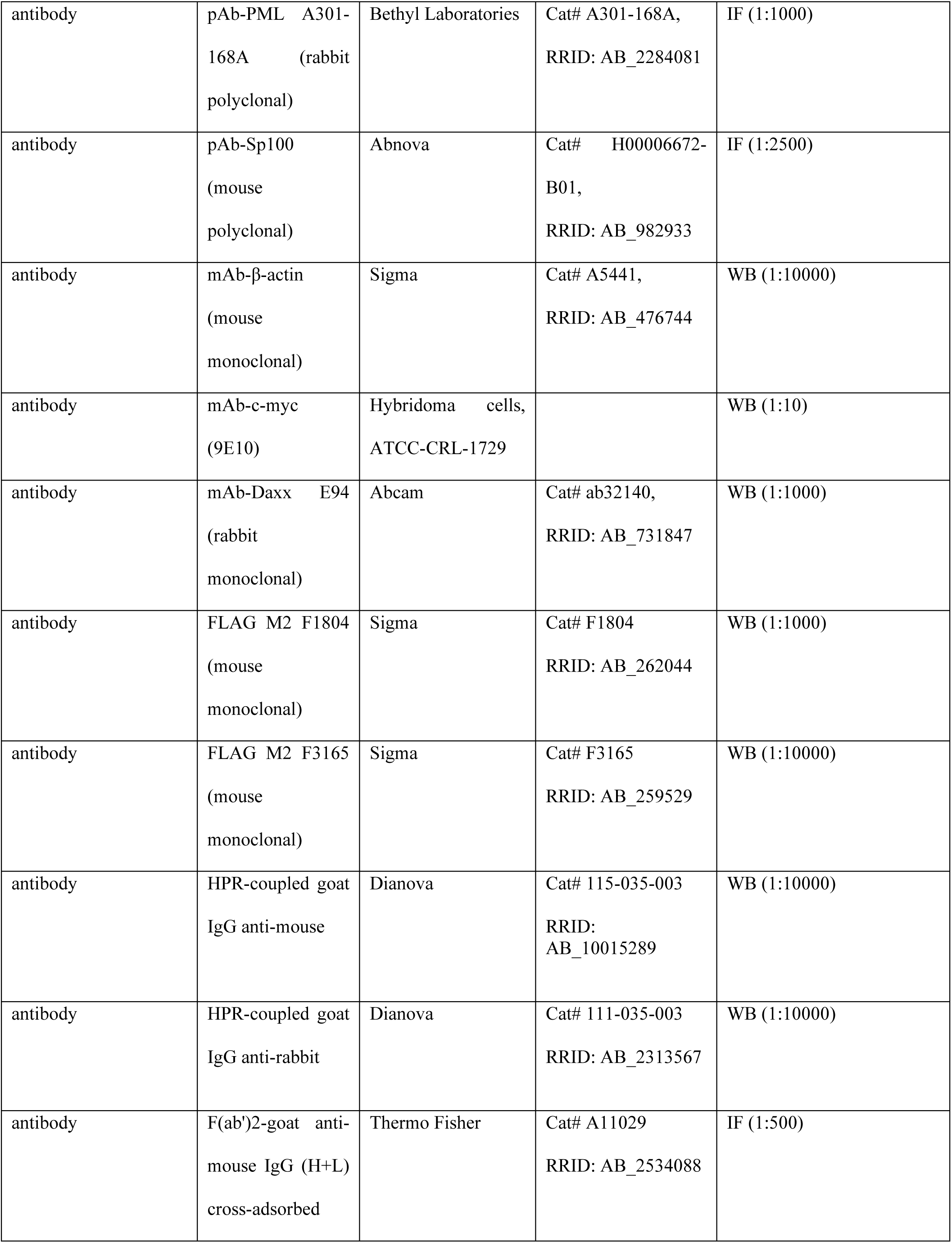
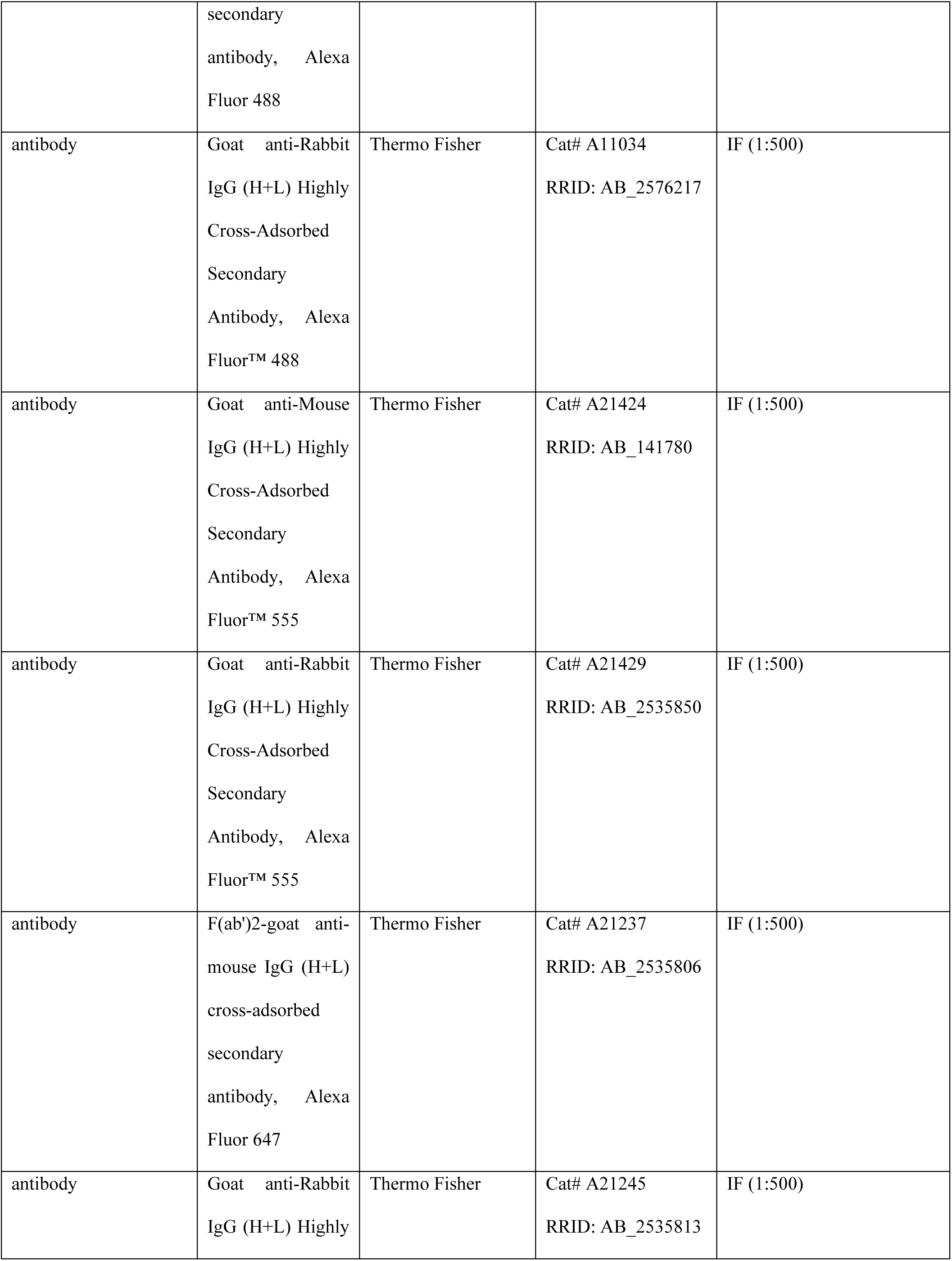
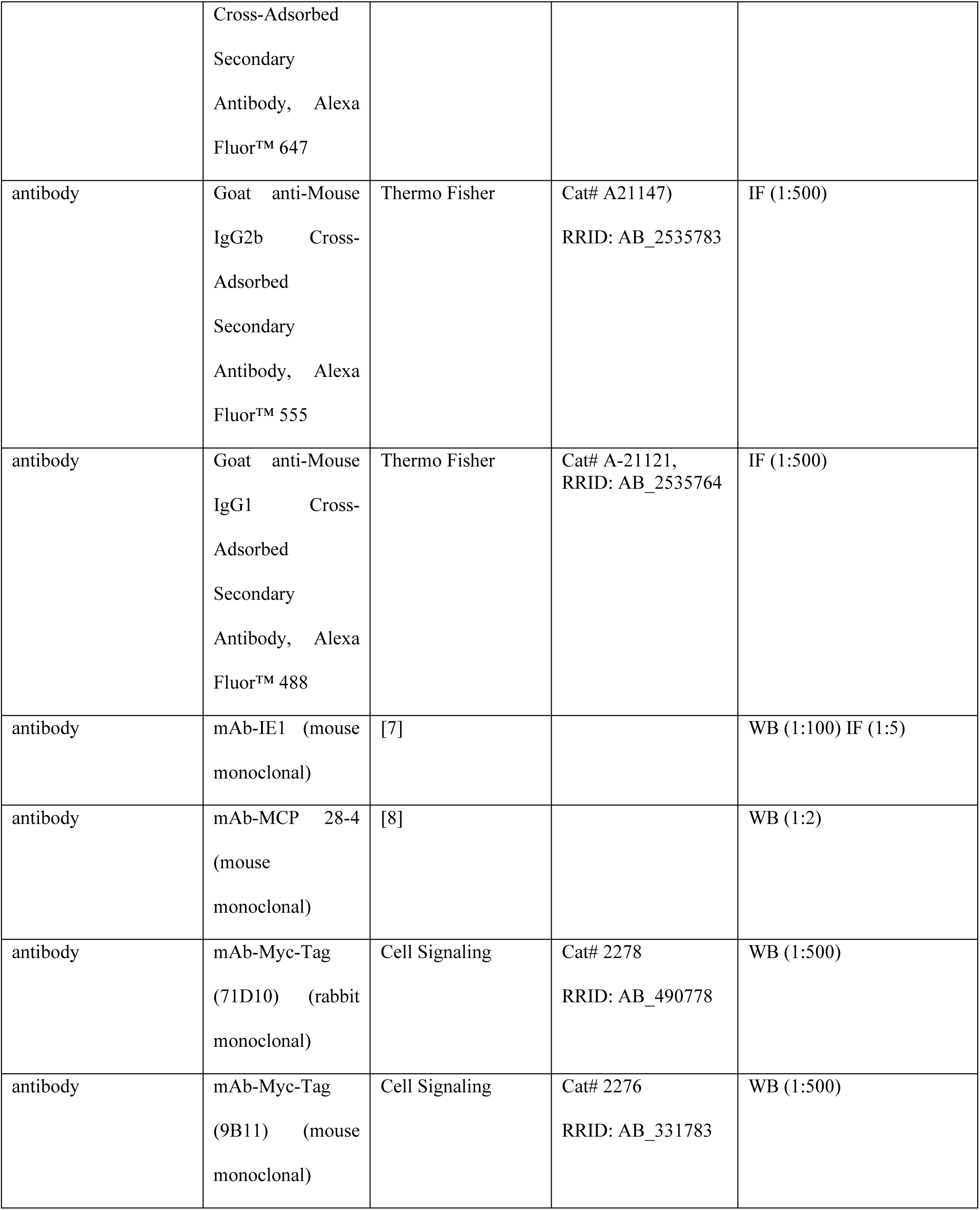
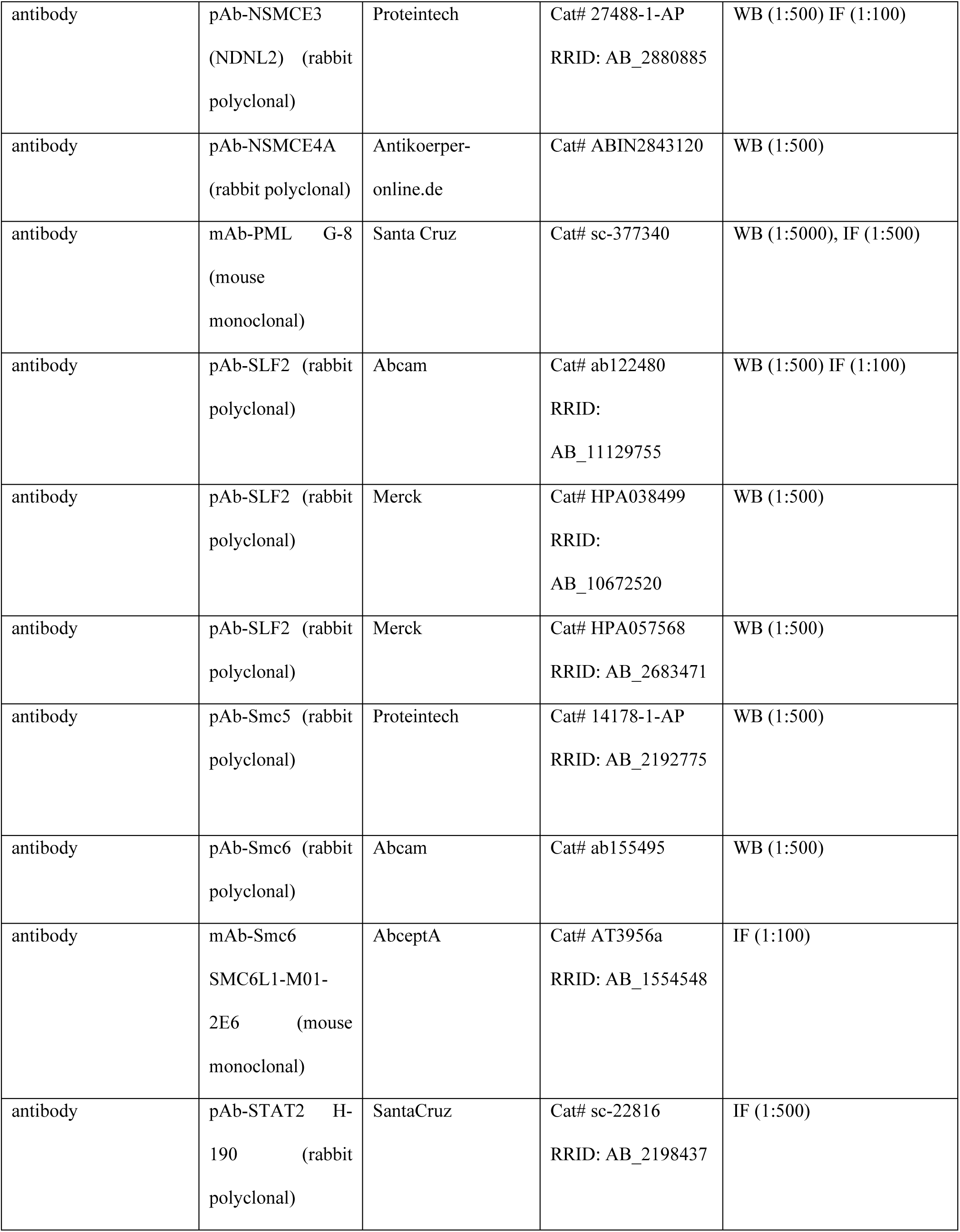
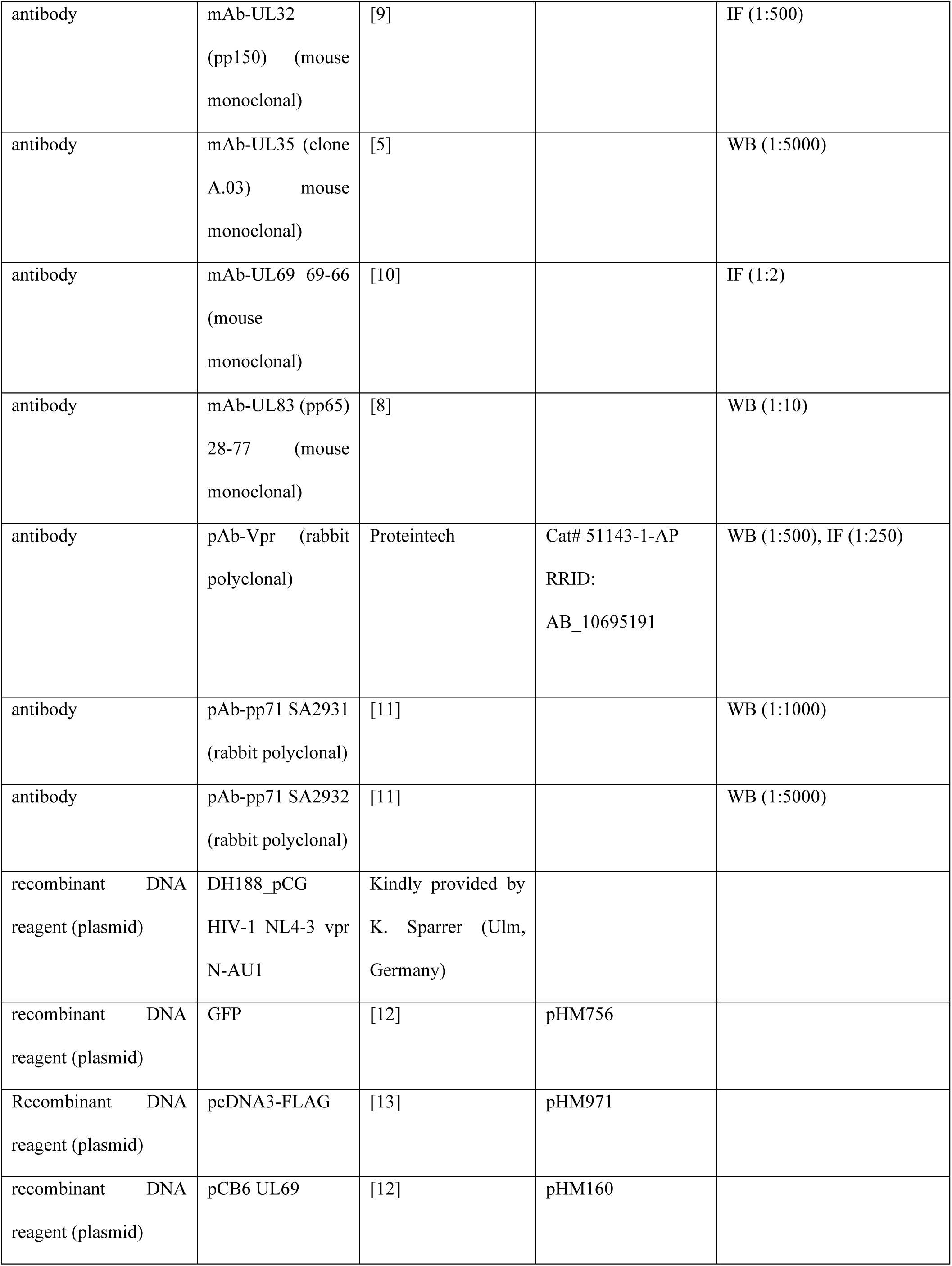
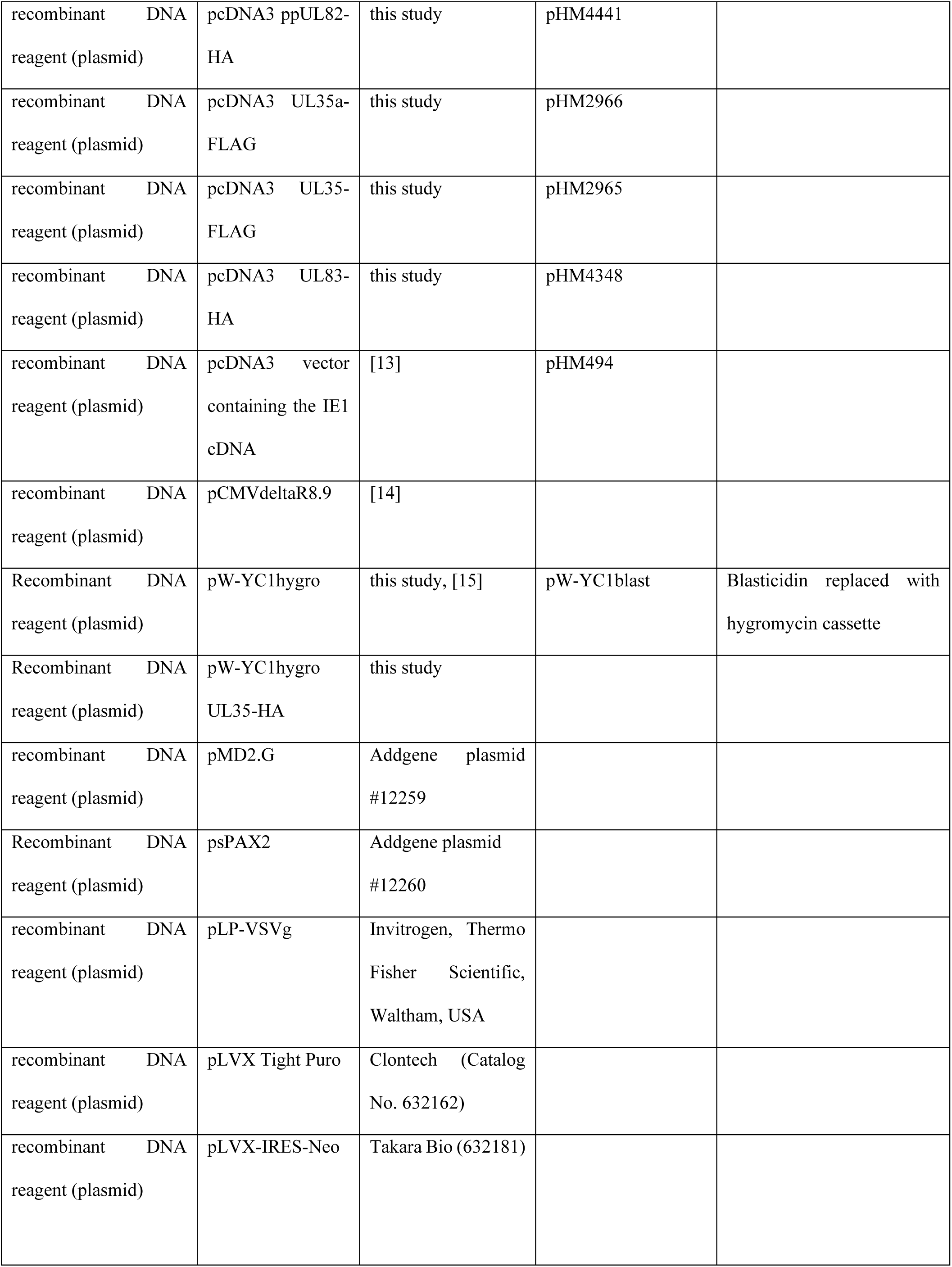
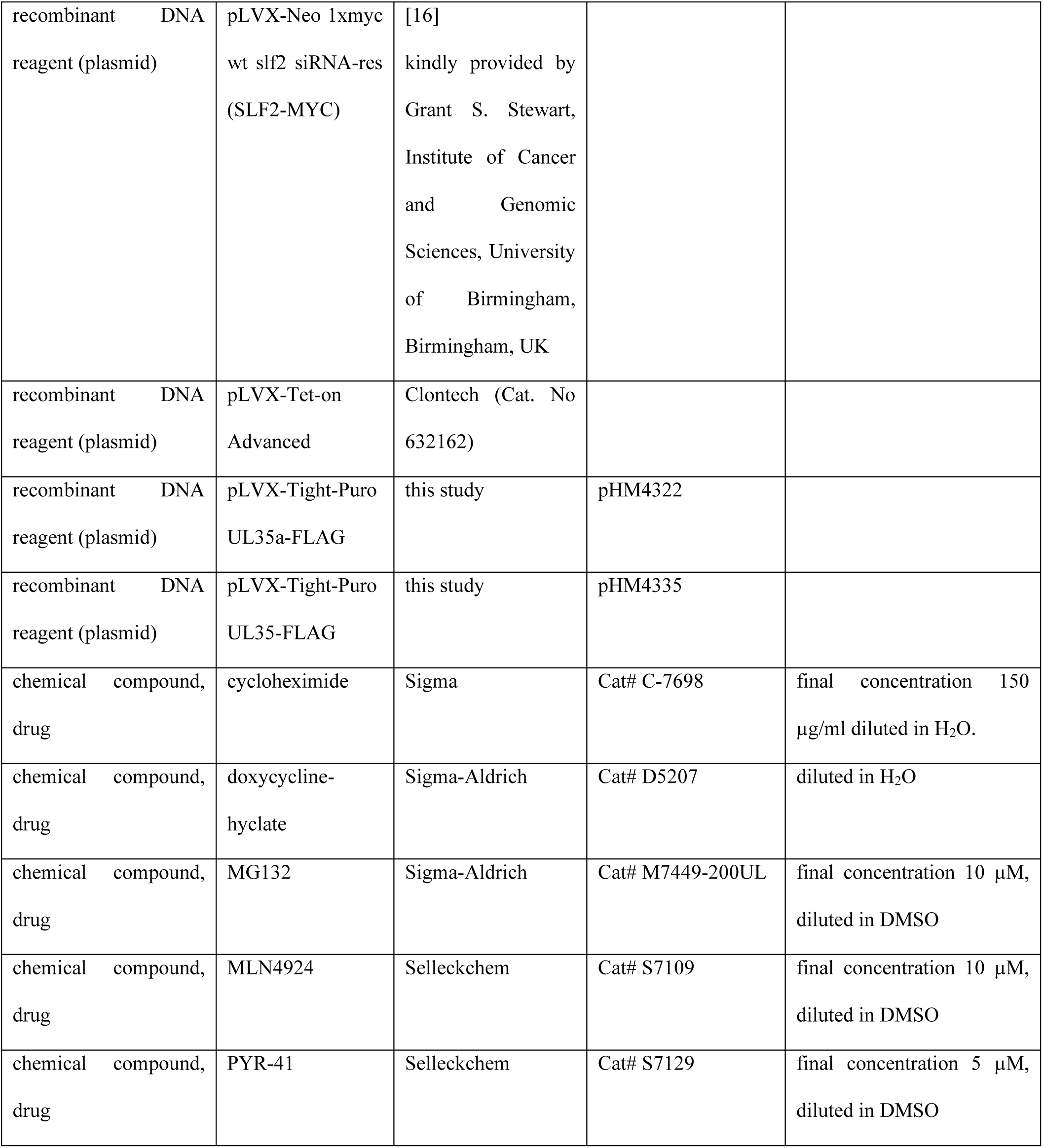
Key resources table.

---

**S2 Table:**
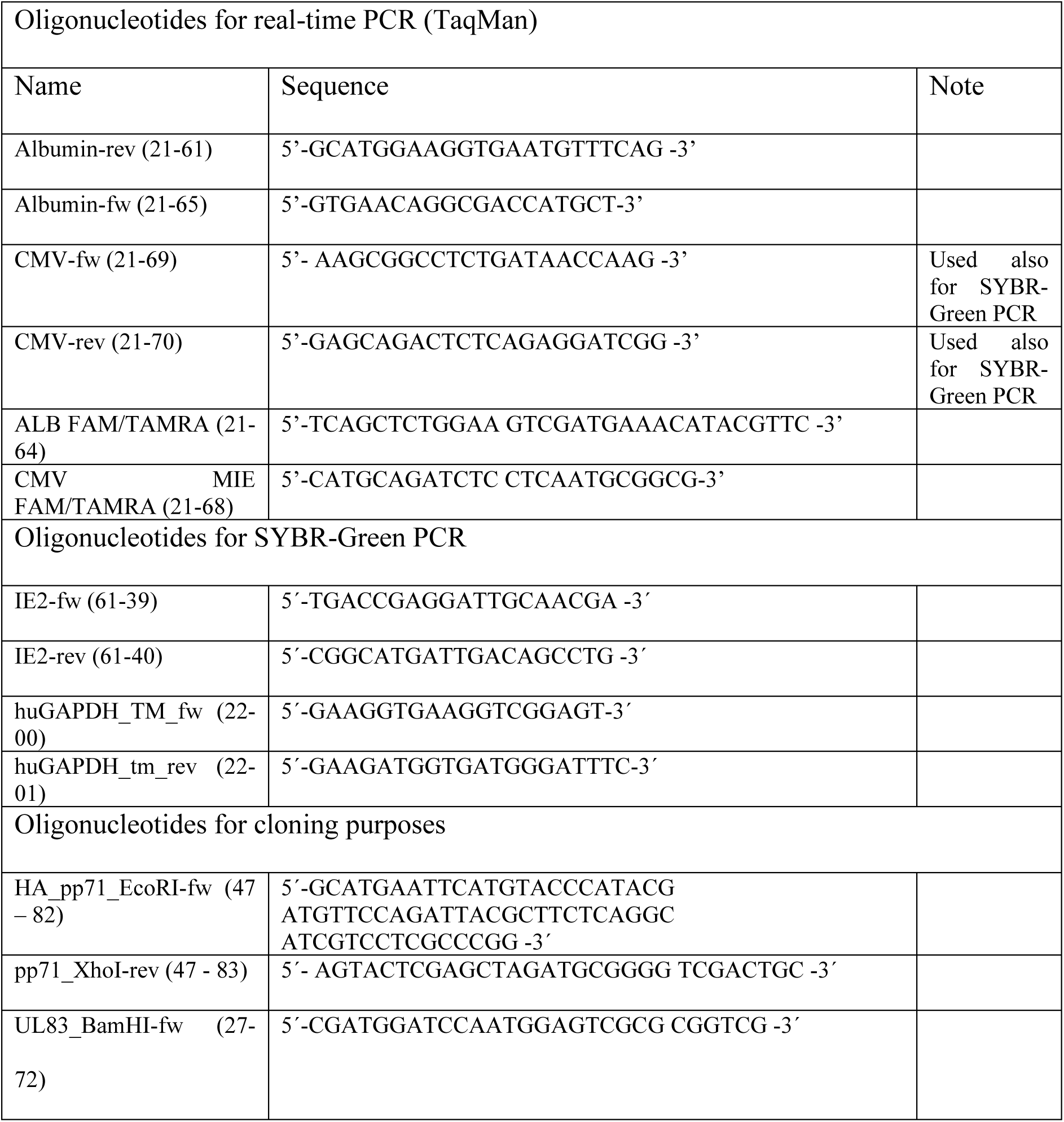
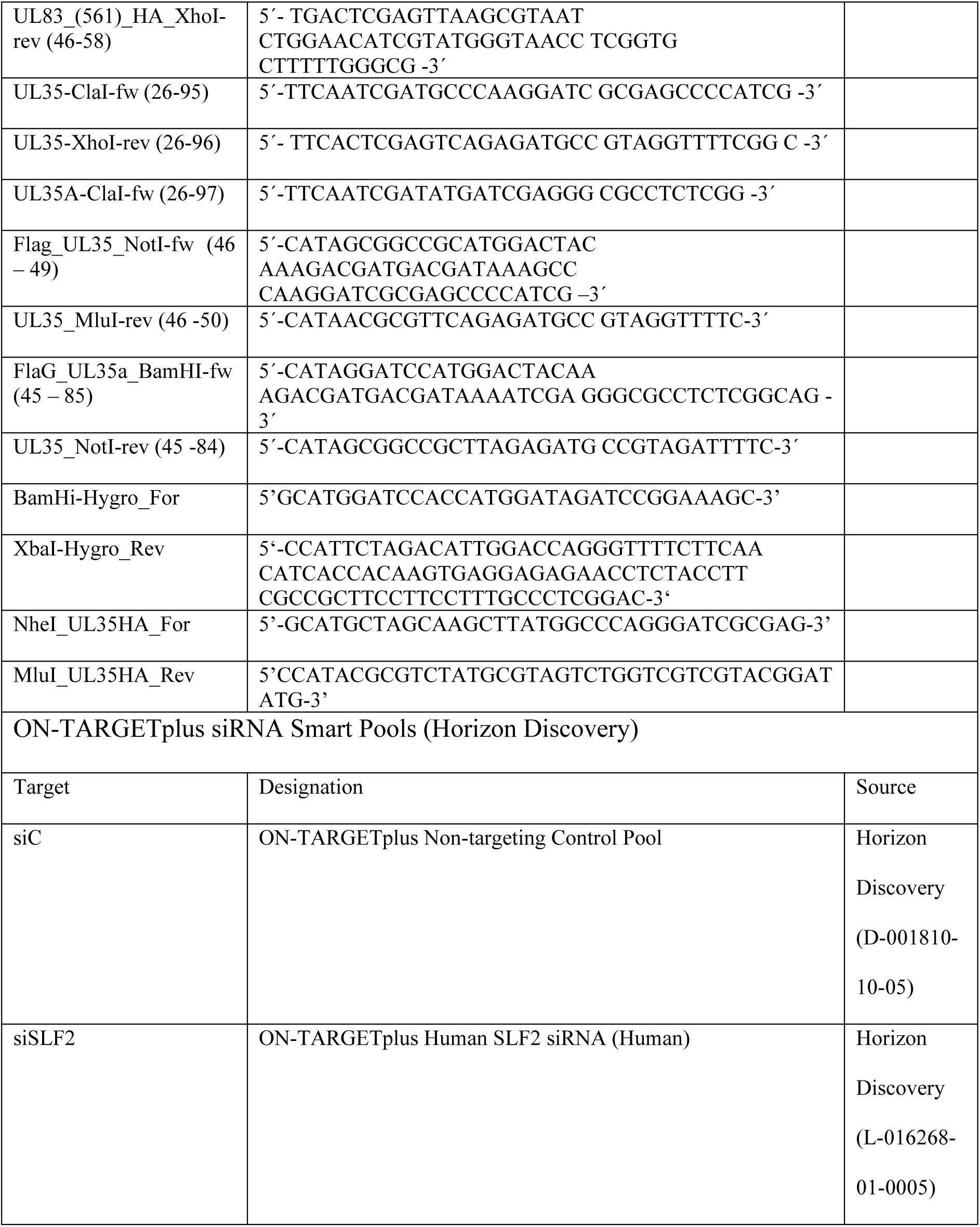
Oligonucleotides.

